# Divergent transcriptional responses of conserved cellular machinery to warming and phosphorus limitation in freshwater phytoplankton

**DOI:** 10.64898/2026.10.28.760975

**Authors:** Joana L. Santos, Vanessa Weber de Melo, Sarah Levasseur, Alexandra A.-T. Weber, Anita Narwani

## Abstract

Lakes are simultaneously warming and becoming phosphorus limited, but whether phytoplankton taxa of the same phylum respond through similar molecular mechanisms remains unclear, challenging a common assumption underlying predictions of ecological responses to environmental change. We cultured four freshwater phytoplankton species, including two diatoms (*Fragilaria crotonensis* and *Nitzschia palea*) and two cyanobacteria (*Microcystis aeruginosa* and *Synechococcus sp.*), across factorial gradients of temperature (20, 24 and 28 °C) and phosphorus (P) limitation, and characterized gene expression using RNA-seq, differential expression analysis and signed gene set enrichment. We generated predictions for expected responses for categories of gene functions derived from the growth rate (GRH) and temperature-dependent physiology hypotheses (TDPH). Warming and P limitation engaged largely the predicted functional categories in all species, however the direction of the response was not conserved, even within phyla—the two cyanobacteria regulated photosynthesis, ribosomal and chaperone genes in opposite directions. Ribosome- and translation-related genes gave the strongest warming response and, in *N. palea*, reversed direction within the species, increasing between 20 and 24 °C and decreasing between 24 and 28 °C, below the strain’s thermal optimum estimated under moderate P limitation. Ribosomal allocation thus supported each hypothesis in different species and temperature ranges and was not predicted by thermal position alone. Where the interaction effects could be tested, P limitation rarely recruited novel pathways but modified existing warming responses, suppressing them in *F. crotonensis* and amplifying and redirecting them in *M. aeruginosa*. Molecular responses to combined drivers therefore diverge among related taxa, cautioning against aggregating species into functional groups when forecasting phytoplankton responses to global change.

## Introduction

Climate change is simultaneously altering temperature and resource availability in freshwater ecosystems, affecting phytoplankton physiology, population dynamics and ecosystem functioning. Rising surface-water temperatures are intensifying and prolonging thermal stratification in many lakes, reducing vertical mixing and the replenishment of nutrients from deeper waters (Kraemer et al., 2021; Yankova et al., 2017). Concurrently, changes in nutrient loading, including intentional reductions associated with lake restoration, are altering the availability of essential resources for primary producers with impacts on their productivity and stoichiometry (Frenken et al., 2023). Phytoplankton play a fundamental role in aquatic ecosystems, supporting pelagic food webs and regulating carbon and nutrient cycling, so understanding how they respond to the combined effects of warming and nutrient limitation is essential for predicting the ecological consequences of environmental change. The independent effects of temperature and resource availability on phytoplankton growth and metabolism are relatively well understood, but their interactive effects remain difficult to predict. Hutchins and Tagliabue (2024) highlighted how warming and nutrient limitation can interact to alter phytoplankton thermal performance, elemental-use efficiencies and nutrient requirements, with consequences for biogeochemical cycling. Because these interactions may depend on the distinct temperature sensitivities and elemental requirements of different cellular processes, linking environmental change to phytoplankton physiology and ecological performance requires understanding responses at the molecular level.

Temperature and nutrient availability regulate phytoplankton growth through fundamentally different mechanisms. Temperature directly influences enzymatic reaction rates, affecting processes such as photosynthesis, respiration and protein synthesis, whereas nutrient limitation constrains metabolic activity by restricting the availability of substrates and essential cellular components (Brown et al., 2004; Sterner & Elser, 2002). Phosphorus (P) is particularly important because it is required for nucleic acid synthesis, ribosome production, membrane phospholipids and cellular energy transfer. The Growth Rate Hypothesis (GRH) proposes that rapidly growing organisms require greater investment in P-rich ribosomal RNA to support increased protein synthesis, linking cellular P requirements to growth and biomass elemental stoichiometry (Elser et al., 2000; Sterner & Elser, 2002). However, the Temperature-Dependent Physiology Hypothesis (TDPH) makes a different prediction, by suggesting that warming can increase the efficiency of ribosomal protein synthesis, allowing phytoplankton to maintain or increase protein production with fewer ribosomes and consequently lower cellular P requirements (Armin & Inomura, 2021; Toseland et al., 2013). These predictions do not apply uniformly across the cell, since cellular components vary in their elemental requirements. Ribosomes contain large quantities of P-rich rRNA, whereas photosynthetic machinery, including photosystem proteins, light-harvesting complexes and carbon-fixation enzymes, requires substantial investment in carbon and nitrogen. Warming may reduce the ribosomal machinery needed to support a given growth rate while also altering the balance of investment among photosynthesis, biosynthesis and cellular maintenance (Armin & Inomura, 2021; Toseland et al., 2013). In contrast, under P-limitation, phytoplankton are expected to reduce P-demanding processes specifically and increase investment in P acquisition and recycling. Consistent with both factors influencing ribosomal investment, Hessen et al. (2017) found higher cellular RNA content at lower temperatures in *Chlamydomonas reinhardtii*, while phosphorus availability strongly influenced RNA content, growth and alkaline phosphatase activity. Warming and P limitation may therefore have partly convergent effects for other cellular functions, though severe P limitation may constrain the synthesis of essential cellular components needed to sustain the higher metabolic activity stimulated by warmer conditions.

Recent studies show that temperature and resource availability jointly influence phytoplankton growth and physiology, often in species-specific ways. Resource limitation can modify or override temperature effects on stoichiometry and metabolism and frequently has stronger effects than temperature on biochemical composition (Levasseur et al., 2025; Weber De Melo, Suter, et al., 2025; Weber De Melo, Thomas, et al., 2025).

Transcriptomic approaches can provide mechanistic insights into phytoplankton responses to environmental change by revealing the coordinated regulation of cellular functions underlying these physiological responses. Under P limitation, phytoplankton increases the expression of genes involved in phosphate transport, phosphorus acquisition and recycling, while downregulating genes associated with P-demanding processes such as nucleic acid synthesis and ribosome biogenesis, and reorganise carbon metabolism and membrane lipid composition (Alipanah et al., 2018; Dyhrman et al., 2012; Harke & Gobler, 2013). Warming also induces substantial transcriptional reorganisation in phytoplankton, affecting photosynthesis, energy metabolism, protein synthesis and cellular stress responses. However, the direction and magnitude of these changes depend on species, exposure duration and temperature relative to their thermal optimum (Hong et al., 2025; Thangaraj & Sun, 2021). Importantly, the transcriptional responses to warming and P limitation may not be independent, as they draw on overlapping cellular machinery. We distinguish three potentially co-occurring forms of interaction (Alipanah et al., 2018; Armin & Inomura, 2021; Hessen et al., 2017; Toseland et al., 2013). Compensatory responses, where warming partly offsets P limitation and weakens P-dependent responses. Constrained responses, where insufficient P prevents or attenuates warming-induced transcriptional changes in biosynthesis, cell division and energy metabolism. Novel responses, where the combined treatment activates pathways not induced by either driver alone, indicating regulatory reorganisation. These mechanisms are not mutually exclusive. We therefore predict that P availability will modify warming responses, with patterns differing among species. Our specific expectations for each functional category are summarised in Table 1.

**Table 1.**
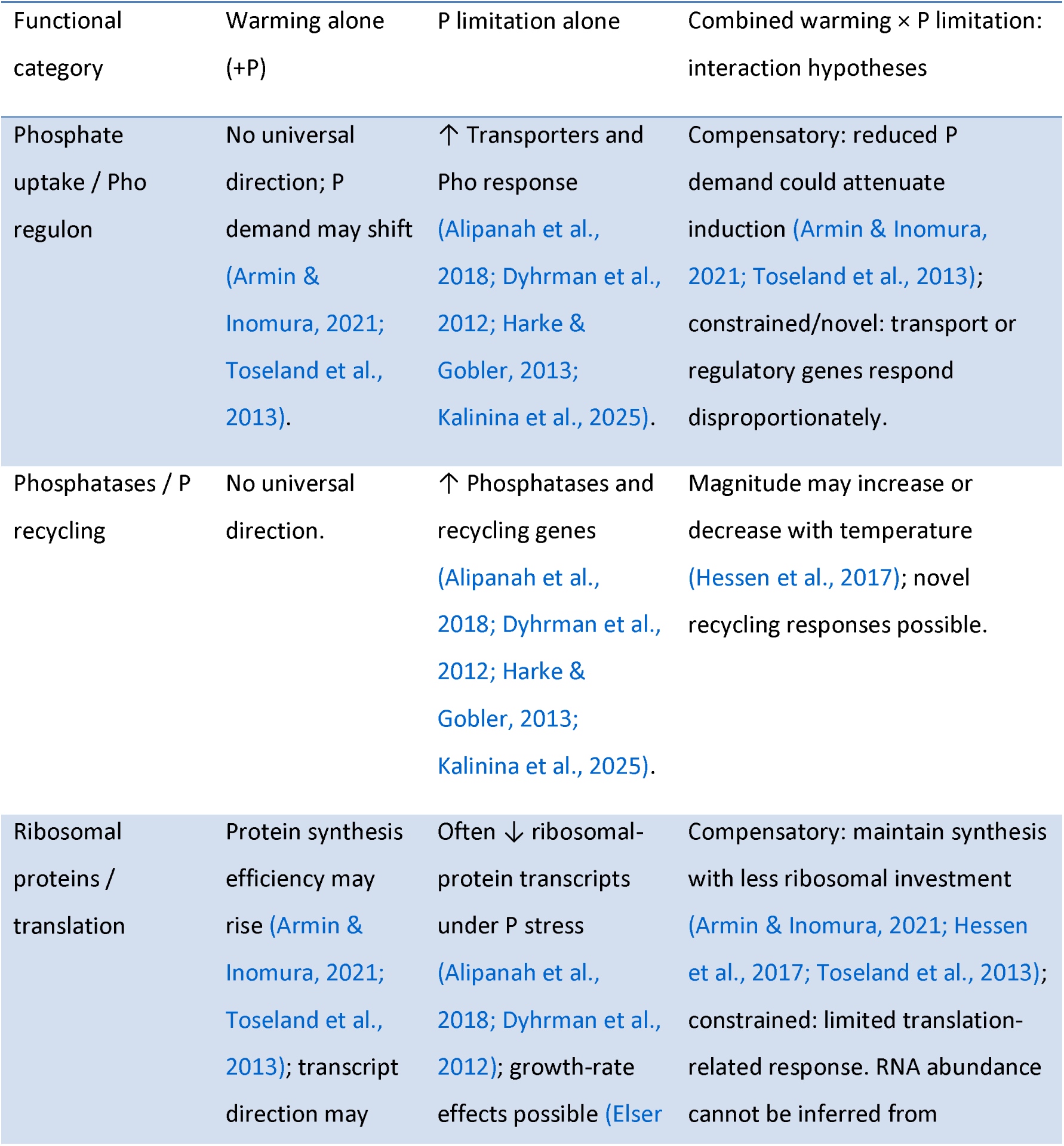
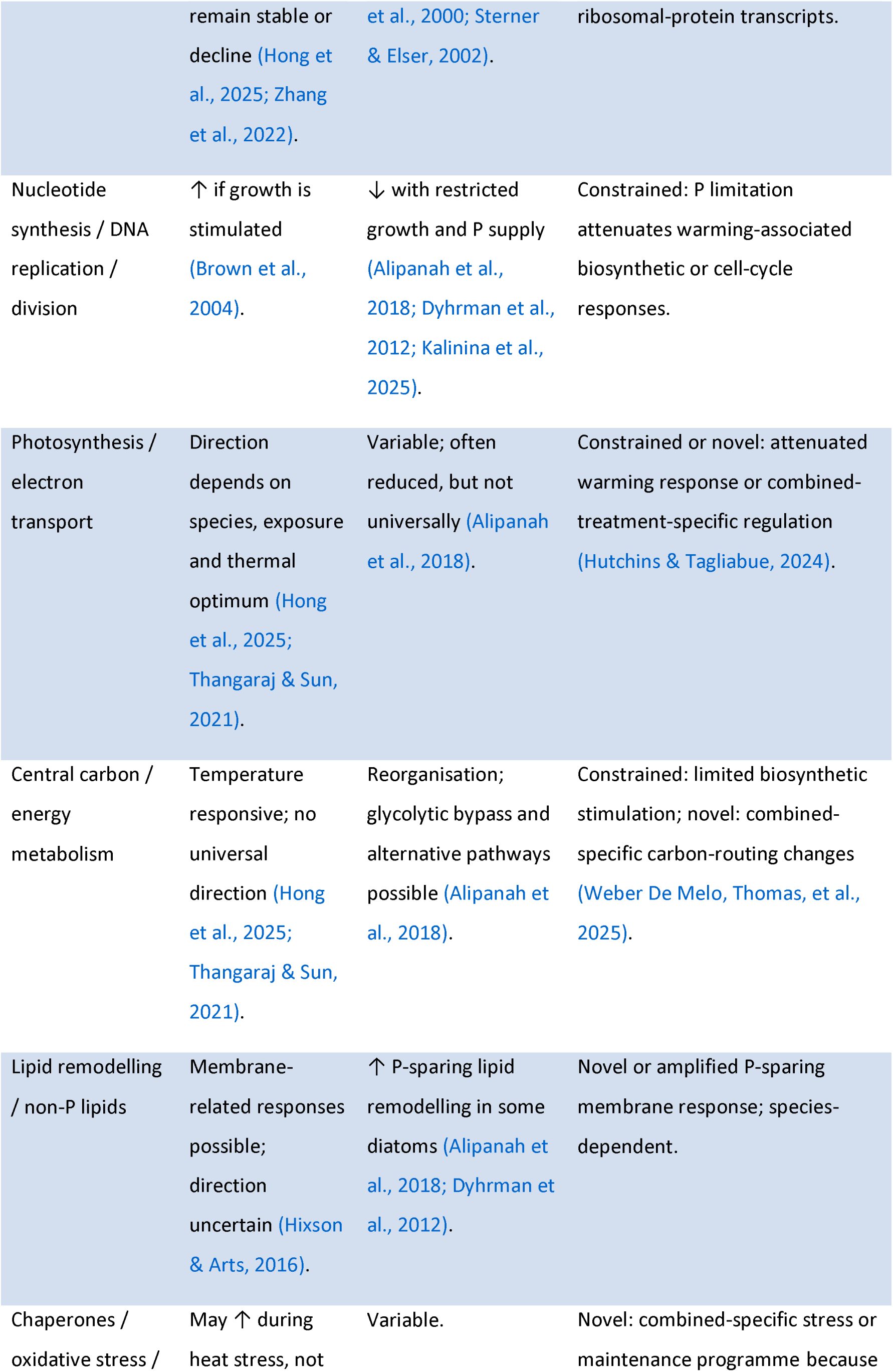
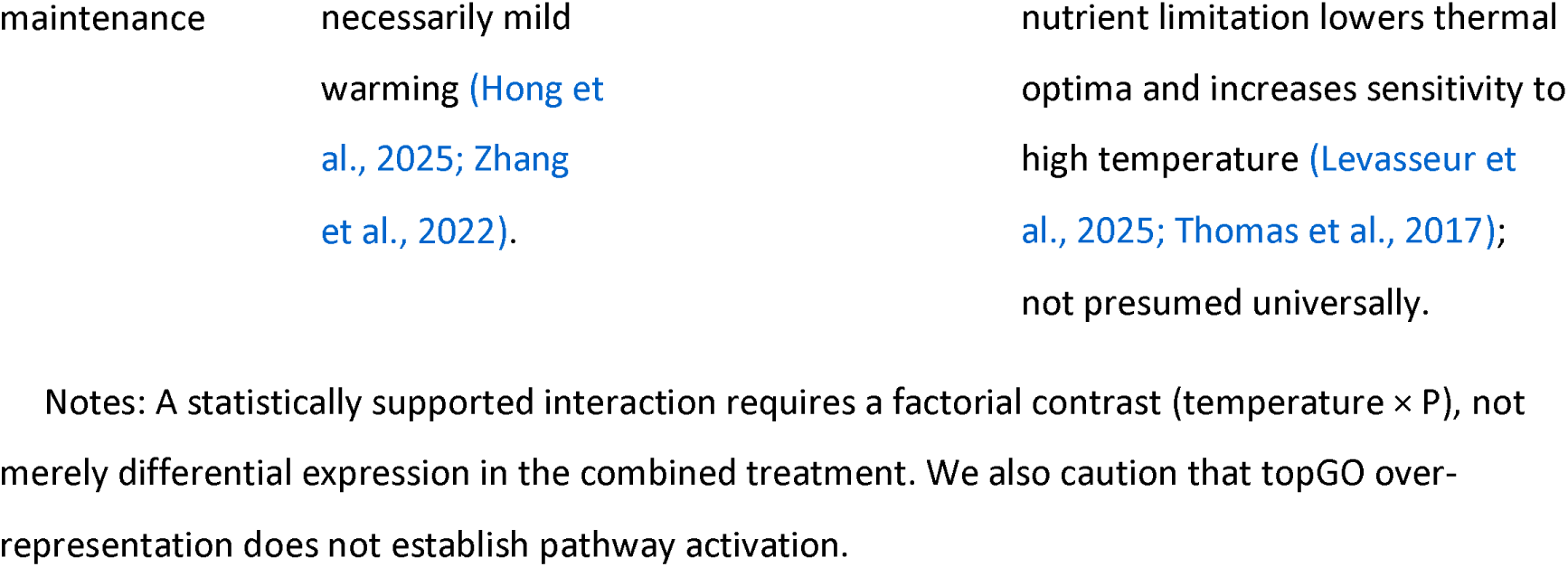
Predicted transcriptional responses of key functional categories to warming, phosphorus (P) limitation and their combination. For the combined treatment, predictions are framed as three interaction types: compensatory (warming makes protein synthesis more efficient, lowering P demand and weakening P-dependent responses), constrained (P limitation prevents or attenuates warming-associated responses) and novel (genes or pathways respond to the combination but weakly or not at all to either driver alone). Arrows give the expected direction of transcript abundance or of directional enrichment (↑, increase; ↓, decrease), not measured pathway flux. Predictions for warming assume temperatures within the physiologically relevant range; responses near or above the thermal optimum may differ.

Here, we investigate how warming and P limitation independently and interactively influence gene expression in four freshwater phytoplankton species: the diatoms *Fragilaria crotonensis* and *Nitzschia palea*, and the cyanobacteria *Synechococcus sp.* and *Microcystis aeruginosa*. Cultures were grown under the temperature (20, 24 and 28 °C) and P limitation treatments (moderate and low) of an accompanying stoichiometry experiment Levasseur *et al*. (accepted), and transcriptomes were analysed for differential expression, functional over-representation and coordinated directional shifts. The interaction effect could be tested in two of the four species, as low biomass prevented extraction of sufficient RNA material in some limiting treatments. We ask: (1) How do warming and P limitation individually affect gene expression and in which direction? (2) does P limitation modify transcriptional responses to warming? and (3) are transcriptional responses shared across species or taxon-specific? By comparing functional responses across four species, we aim to connect stoichiometric theory to the molecular mechanisms underlying phytoplankton responses to multiple environmental drivers.

## Materials and Methods

### Study system and species

We quantified gene expression dynamics in four freshwater phytoplankton species. Three species were previously isolated from lakes Greifen and Constance in northeastern Switzerland, while *Synechococcus* sp. (CCAP 1478/11) was obtained from the Culture Collection of Algae and Protozoa. The species represent two major phytoplankton groups: Bacillariophyta (*Fragilaria crotonensis*, *Nitzschia palea*) and Cyanobacteria (*Microcystis aeruginosa*, *Synechococcus* sp.). Detailed information on species isolation, identification, and culture history is provided in (Levasseur et al., 2025). The strains used here correspond to *N. palea* C and *M. aeruginosa* strain 1 from that study. Cultures were non-axenic but maintained in a mineral medium (COMBO, (Kilham et al., 1998)) without organic carbon to minimize bacterial growth. Prior to the experiments, all species were acclimated in batch culture at 15°C under nutrient-replete conditions, with a 14L:10D light cycle and low irradiance (20 μmol photons m^-2^s^-1^).

### Experimental design and growth conditions

Samples came from the phosphorus (P) experiment of a larger study of phytoplankton stoichiometry that manipulated nitrogen, P and light separately (Levasseur et al., accepted), Each species received six treatments (Fig.1a), each with three biological replicates. Three temperatures (20, 24 and 28 °C) were crossed with two levels of P limitation, moderate (P-mod, 10 µmol L⁻¹ K₂HPO₄) and severe (P-low, 0.1 µmol L⁻¹), with all other resources supplied in excess. At P-mod, the thermal optima of these strains are 23.9 °C for *F. crotonensis*, 32.2 °C for *N. palea* and 26.1 °C for *Synechococcus sp*.; at P-low, an estimate exists only for *N. palea* (16.2 °C, 95% credible interval (CI) 1.6–25.0 °C). No optimum could be estimated for *M. aeruginosa* strain 1, whose growth increased up to the highest temperature tested (35 °C) (Levasseur et al., 2025).

Populations were acclimated first to temperature, raised by 2 °C per day and then held for 48 h at the target temperature, and then to P, by centrifugation and resuspension in medium with the target P concentration (Fig. 1b; Appendix S1). Cultures were grown in sterile, ventilated 500 mL tissue-culture flasks, inoculated at a standardised low biomass (20 RFU of pigment fluorescence in 350 mL of COMBO medium), shaken at 70 rpm under an 18L:6D light cycle, and monitored daily by fluorescence until growth approached steady state or for at most ten days (Fig. 1b, Appendix S1). See Levasseur *et al*. (accepted) for further experimental details.

**Figure 1.**
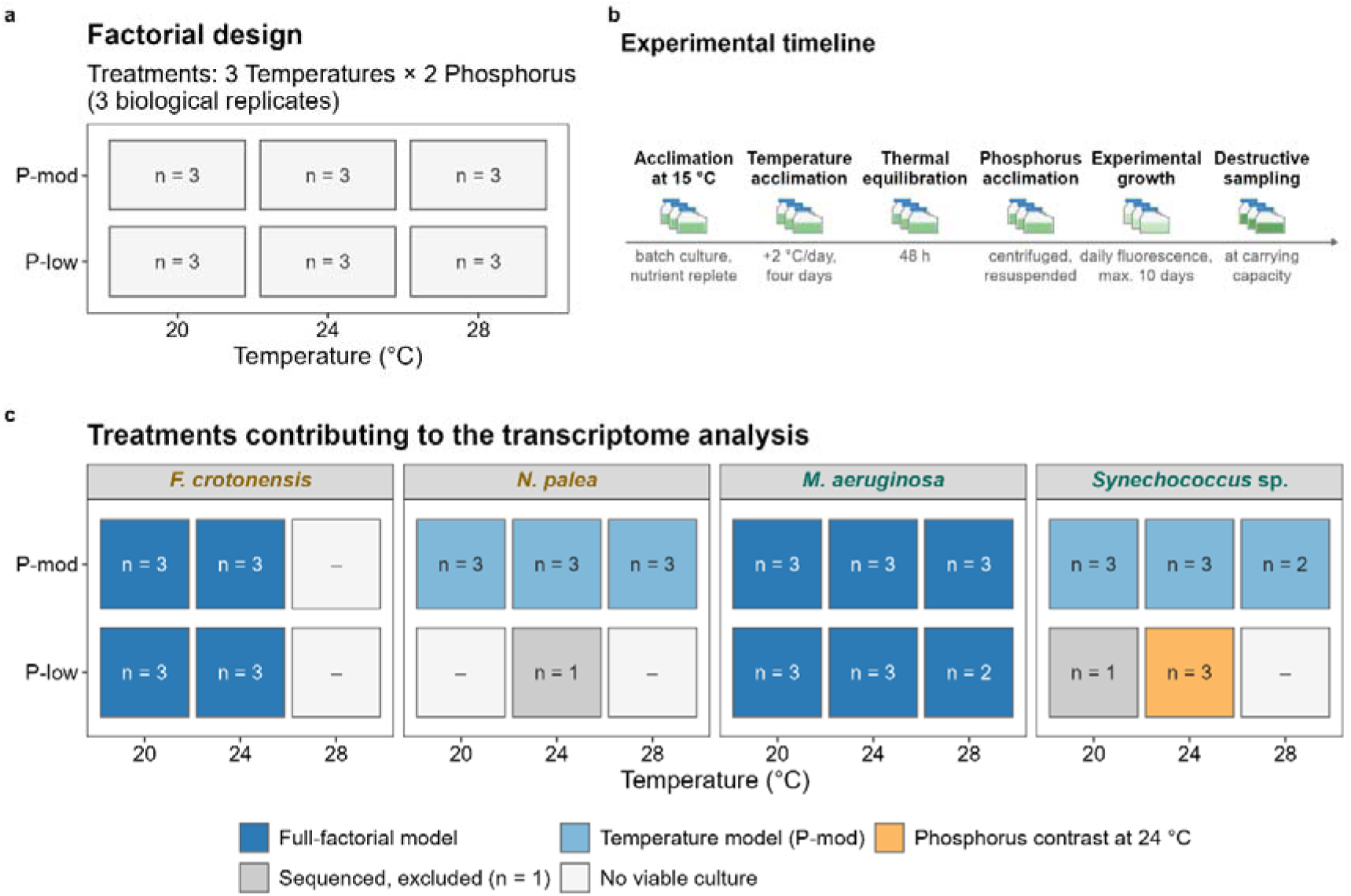
Experimental design and treatments included in the transcriptomic analysis. (a) Planned full-factorial design, with three temperatures (20, 24 and 28 °C) crossed with two phosphorus (P) availabilities, moderate (P-mod, 10 µmol L⁻¹ K₂HPO₄) and limiting (P-low, 0.1 µmol L⁻¹) for each species. Each treatment started with three biological replicates. (b) Experimental timeline showing the different stages and their duration. (c) Realised design after accounting for treatment-specific culture failure and low RNA yield. Numbers indicate sequenced biological replicates. Colours show the analysis in which each treatment was included: full-factorial model (dark blue), temperature series under P-mod (light blue), and P contrast at 24 °C (orange). Treatments with a single replicate (grey) were included in ordination analyses but not in differential expression models; –, no viable culture. Species names are colour-coded throughout figures, with diatoms in golden brown and cyanobacteria in green.

### Transcriptome profiling

Experimental populations were monitored until steady-state and were then sampled destructively to obtain biomass for gene expression analysis (Fig. 1b). Only combinations of temperature and P treatments where more than one replicate produced adequately high quantities of good quality mRNA for sequencing were used for the following gene expression analysis (Table S1). The treatments retained for each species are given in Fig. 1c. RNA extraction was done with the E.Z.N.A HP Total RNA kit, Omega Biotek. Library preparation was performed with the Illumina Stranded Total RNA Prep kit for the cyanobacteria and the Illumina Stranded mRNA Prep. kit for the diatoms, following manufacturer instructions. Libraries were sequenced on an Illumina NovaSeq X by the Functional Genomics Center Zurich.

### Differentially expressed gene (DEG) analysis

To ensure high-quality sequences, rRNA and adapters were removed before transcriptome mapping using bbmap v.39.05 (Bushnell, 2014) (k=21 and minlen=100) and fastp v.0.23.2 (S. Chen et al., 2018) (-l 100 and disable-quality-filtering). Paired-end RNA-seq reads were pseudoaligned and quantified with kallisto v.0.46 (Bray et al., 2016) (default setting including - single-overhang) using transcripts from the reference genomes of *F. crotonensis* (GCA_022925895.1), *N. palea* (GJPI00000000.1), *M. viridis* (GCF_014698335.1) for *M. aeruginosa*, and *S. elongatus* (GCF_019038515.1) for *Synechococcus sp.*, as no annotated genome exists for the two cyanobacteria strains. After rRNA and adapter removal, libraries contained on average 34.0 ± 7.7 million read pairs in *F. crotonensis*, 25.0 ± 4.8 in *N. palea*, 29.8 ± 15.8 in *M. aeruginosa* and 24.0 ± 9.1 in *Synechococcus* sp. (mean ± SD), of which 29.4 ± 2.9%, 29.5 ± 8.2%, 49.1 ± 5.7% and 46.1 ± 20.3%, respectively, pseudoaligned to the reference transcripts (Table S1). Transcript abundances were imported into R v4.4 (R Core Team, 2024) using tximport v1.34.0 (Soneson et al., 2015) (type=”kallisto”, txOut=TRUE), retaining transcript-level estimates.

Differential expression was analysed with edgeR v4.4.2 (Y. Chen et al., 2025) correcting for both library size and transcript-length bias. Library sizes were normalised with the trimmed mean of M-values (TMM) method implemented in calcNormFactors (Y. Chen et al., 2025; Robinson et al., 2010), and combined with transcript lengths into a single offset supplied through scaleOffset (Appendix S1). Lowly expressed genes were removed with filterByExpr using the design matrix, following the treatment structure. Negative binomial dispersion parameters were estimated with estimateDisp, and a quasi-likelihood negative binomial generalised linear model was fitted with glmQLFit.

Two complementary differential expression tests were performed for each species. Model level effects were tested using glmQLFTest and summarized with topTags, using all coefficients for the temperature and P factors and their interaction in a single full-factorial design model (∼Temperature x P). Pairwise contrasts approach tested specific comparisons defined by makeContrasts from the limma v3.62.2 R package (Law et al., 2018; Ritchie et al., 2015), derived from the model. Temperature contrasts evaluated each level (24 vs 20, 28 vs 20 and 28 vs 24 °C) within each phosphorus level, and P contrasts evaluated P-low relative to P-mod within each temperature, where the experimental design allowed. The P contrasts were defined so that the tested level is P-low, matching the temperature contrasts in which the tested level is the warmer temperature; a positive log2 fold change or normalised enrichment score therefore indicates higher expression under phosphorus limitation. The full-factorial model could be fitted for *F. crotonensis* and *M. aeruginosa* (Fig. 1c). For *N. palea* and *Synechococcus* sp., for which a full temperature series was only available at P-mod, we analysed a restricted model (∼Temperature), which does not allow testing for interactive effects, but does allow us to explore certain pairwise contrasts. Specifically, for *Synechococcus sp.* we used an independent model to analyse the contrast of P-low versus P-mod at 24 °C.

For main model effects, genes were considered differentially expressed at a Benjamini–Hochberg false discovery rate (FDR) < 0.05. For pairwise contrasts, which provide directly interpretable log2 fold changes (log2FC) from linear combinations of model coefficients, significance required both FDR < 0.05 and |log2FC| ≥ 0.5. Significant DEGs from pairwise contrasts were classified as up- or downregulated based on the sign of the log2FC.

Transcriptome-wide variation was summarized per species by principal component analysis and multidimensional scaling of log_2_ counts per million, using all samples of a species, including those not entering the models (Appendix S1).

### Gene ontology enrichment analysis

Functional annotation combined the published genome annotations, available for the two cyanobacteria reference genomes, with InterProScan predictions generated for the predicted proteomes of all four species. InterProScan was the main source of annotation and the only one for *F. crotonensis* and *N. palea*. Protein sequences were extracted from each genome and its corresponding annotation using gffread v.0.12.7 (Pertea & Pertea, 2020), and annotated with InterProScan v5.77-108.0 (Blum et al., 2025; Jones et al., 2014) against all member databases, with InterPro entry and GO term mapping enabled (-iprlookup, - goterms). GO terms from both sources were merged into a single gene-to-GO table per species. Because count matrices, GFF features and InterProScan output do not share a single identifier space, a species-specific mapping step was applied before annotation, in which GFF files were imported with rtracklayer v1.66.0 (Lawrence et al., 2009). The GO enrichment background was defined as the set of transcripts retained by filterByExpr in edgeR v4.4.2 (Y. Chen et al., 2025) and with a valid GO annotation.

GO over-representation was tested with topGO v2.58.0 (Alexa & Rahnenfuhrer, 2024) using the weight01 algorithm, Fisher’s exact test and minimum node size of 5. Each Gene Ontology domain (biological process, BP; molecular function, MF; cellular component, CC) was tested separately for each model effect and pairwise contrast, against a universe of all expressed, GO-annotated genes. For each effect, resulting term-level Fisher *p*-values were adjusted for multiple testing within that effect and within each ontology separately using the Benjamini-Hochberg (BH) procedure, giving both a nominal Fisher *p*-value and FDR-corrected value per GO term (Table S3). GO terms were considered enriched at Fisher *p*<0.05 for an exploratory approach, and separately at FDR<0.05 for a more conservative approach. Nominal Fisher *p-*values are used for the main figures and FDR-significant terms are reported separately (Appendix S1).

Over-representation was tested on all differentially expressed genes together rather than separately for up- and down-regulated genes, which would reduce power. Coordinated directional change was assessed by GSEA (below). To describe the composition of each enriched category, we counted how many of its differentially expressed genes increased and decreased in expression (Appendix S1). To compare species at a common functional level, each significant GO term was mapped to its ancestors within its own ontology using GO.db v3.20.0 (Carlson, 2017), accessed through AnnotationDbi v1.68.0 (Pagès et al., 2024), and matched against the generic GO slim subset (Gene Ontology Consortium, 2023; accessed 26.07.2026). When a term mapped to several slim categories, only the most specific was retained, discarding any slim ancestor that was itself an ancestor of another match. The three ontology roots (biological process, molecular function and cellular component) and the two broadest biological process categories (metabolic and cellular process) were excluded from display, because they would apply to most terms. Enrichment was tested for individual GO terms and the displayed categories carry no *p*-value.

Annotation coverage was assessed to distinguish an absent response from lack of annotation. For gene sets central to our predictions – ribosome and translation, phosphate transport, phosphatases, cell cycle and DNA replication, oxidative stress, membrane lipids, and photosynthesis as a control – the seed GO terms were expanded to include all their descendants (GO.db) and the number of expressed genes carrying any term in each set was counted per species (Table S2). A set containing fewer genes than the minimum node size cannot be tested by either enrichment method. Data were handled with the tidyverse (Wickham et al., 2019), and figures were produced with ggplot2 v4.0.3 (Wickham, 2016), patchwork v1.3.2 (Pedersen, 2025) and ggtext v0.1.2 (Wilke & Wiernik, 2022).

### Gene set enrichment analysis (GSEA)

To complement the previous analyses, a threshold-free approach was applied to the contrast-based results. GSEA was performed using the GSEA function of clusterProfiler v4.14.6 (Xu et al., 2024), which estimated permutation-based p-values with fgsea v1.32.4 (Korotkevich et al., 2019) using a fixed random seed. For each contrast, a ranked gene list was built from the mean log_2_ fold change per annotated transcript. GO terms of all three ontologies and their gene members were taken from the same gene-to-GO tables as for the over-representation analysis. GSEA was run per contrast with gene set sizes restricted to 5-500 genes, a BH correction of *p*-values, and by excluding contrasts with fewer than 10 ranked genes. Significant gene sets (adjusted *p*-values < 0.05) were retained. Each significant gene set is reported with its normalised enrichment score (NES) and adjusted *p-*value in Table S4. Categories containing both positively and negatively enriched gene sets are flagged, since averaging a signed statistic would otherwise mask opposing signals. Because GSEA requires a signed ranking, it cannot be applied to the model-level F-tests. To obtain a directional view of the temperature × P interaction to use as GSEA input, each interaction coefficient was additionally tested individually with glmQLFTest: one coefficient for *F. crotonensis*, and two for *M. aeruginosa* corresponding to 24 °C × P-mod and 28 °C × P-mod.

### Cross-species comparison

For analysis comparing species, GO terms from the over-representation and GSEA analyses were pooled and the summarising step applied once to the combined set, so that both analyses are displayed on an identical set of categories. Within each category, over-representation is summarised as the mean −log₁₀ Fisher p-value and the number of contributing GO terms, and GSEA as the mean NES and the number of contributing terms. To determine whether the same genes responded to more than one effect, the differentially expressed gene sets for temperature, phosphorus and their interaction were intersected within each species, and the size of every intersection was counted. The same procedure was applied to the gene sets recovered by each pairwise contrast. Gene identifiers are not comparable between species, so all intersections are within a species. To test whether phosphorus supply altered the response to warming, the same temperature steps relative to 20 °C were compared at both phosphorus levels in the two full-factorial species, both as the number of genes responding and as the functional categories recovered.

Because the figures display collapsed GO slim categories, which cannot separate specific gene sets, every significant term is reported individually in the Tables S3 (over-representation) and S4 (GSEA), with the category it was assigned to, its ontology and its statistics. Of the 818 significant term-level results, 197 (24%) have no ancestor in the generic GO slim and therefore appear only in these tables.

## Results

### Species-specific transcriptional responses to temperature and phosphorus

Samples clustered by treatment with the dominant source of transcriptional variation differing among the four species (Fig. 2a-d). In *F. crotonensis*, P limitation explained most of the variation, separating P-low from P-mod cultures along PC1 (46.9%; Fig. 2a). In *N. palea*, temperature was the primary driver of variation, with the three temperature treatments ordered along PC1 (27.7%; Fig. 2b). *Microcystis aeruginosa* showed weaker treatment separation, with both environmental drivers contributing to transcriptomic variation (Fig. 2c), whereas *Synechococcus* sp. was structured primarily by P limitation (Fig. 2d). Multidimensional scaling (MDS) gave similar patterns (Fig. S1).

**Figure 2.**
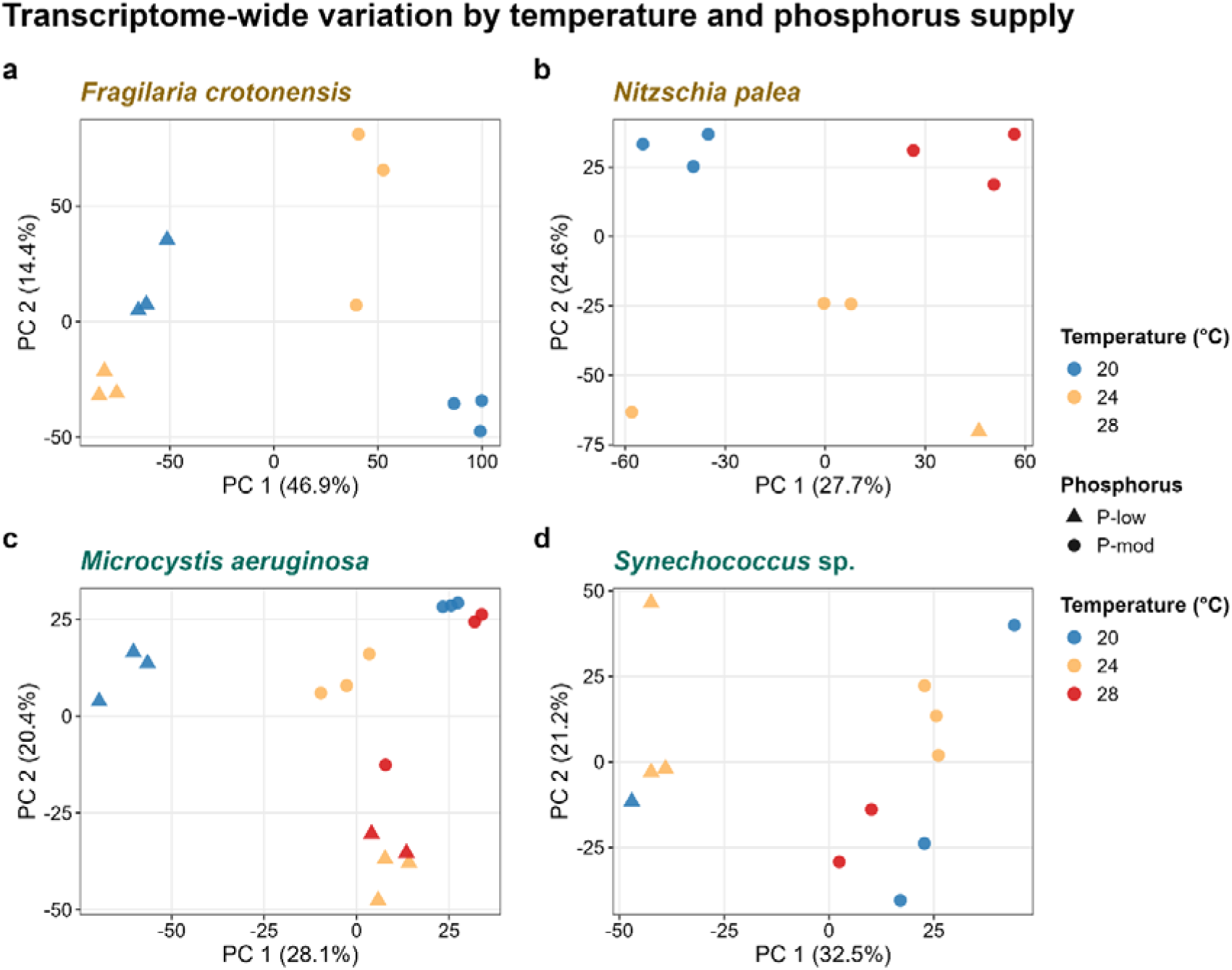
Transcriptome-wide variation among treatments. Principal component analysis of scaled log₂ counts per million for (a) *Fragilaria crotonensis*, (b) *Nitzschia palea*, (c) *Microcystis aeruginosa* and (d) *Synechococcus* sp. Colours show temperature and symbols show phosphorus (P) availability (circles, moderate P (P-mod); triangles, P-low). Axis labels show the proportion of variance explained. All sequenced samples are shown, including treatments excluded from differential expression analyses because only a single replicate was available.

Differential gene expression analyses mirrored the ordination patterns (Fig. 3a). In *F. crotonensis*, P limitation affected a larger fraction of expressed genes (56.4%) than temperature (40.2%) or their interaction (24.2%). The reverse pattern was observed in *M. aeruginosa*, with temperature showing the largest model effect (73.3% of expressed genes), followed by P limitation (57.5%) and the interaction term (51.1%). For both species, P limitation generated the largest numbers of differentially expressed genes, particularly at 20 °C (Fig. S2e). Warming altered the expression of 42.5% of expressed genes in *N. palea*, while *Synechococcus* sp. showed the weakest response to temperature (9.4% of expressed genes; Fig. 3a). In contrast, P limitation produced a stronger response in *Synechococcus* sp., affecting more genes than all temperature contrasts combined (Fig. 3a and S2e).

**Figure 3.**
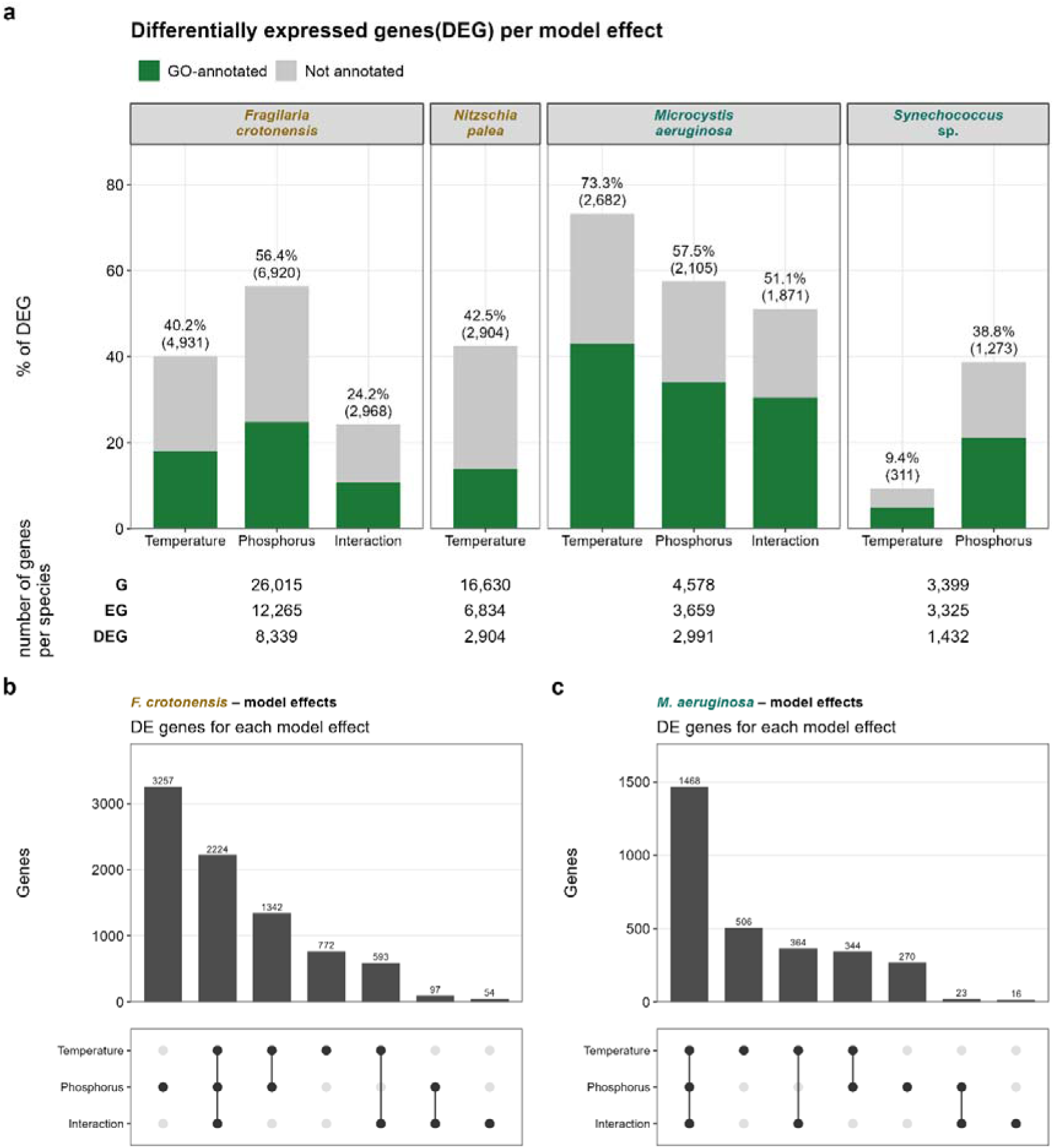
Extent and overlap of differential expression. (a) Differentially expressed genes (DEG) per species and for each model effect, expressed as a percentage of expressed gene (EG). Dark shading indicates genes associated with at least one Gene Ontology (GO) term. Labels give the percentage and numbers of DEGs. (b) The table reports, for each species, the number of genes in the reference annotation (G), genes retained after expression filter (EG; edgeR filterByExpr), and genes differentially expressed for at least one effect (DEG). Because genes may respond to multiple effects, total DEGs do not equal the sum across effects. (b, c) Overlap between the sets of DEG for temperature, phosphorus and their interaction in (b) *F. crotonensis* and (c) *M. aeruginosa*, the two species with a full-factorial design. Bars indicate the number of genes in each intersection; filled dots indicate the corresponding effects. DEGs were identified at FDR < 0.05.

### Extensive overlap in transcriptional responses

Genes responding to temperature, P limitation and their interaction overlapped extensively in the two species, *F. crotonensis* and *M. aeruginosa*, for which a full factorial analysis was possible (Fig. 3b, c). In *F. crotonensis*, 26.7% of differentially expressed genes were shared among all three effects, while this proportion increased to 49.1% in *M. aeruginosa*. By contrast, differentially expressed genes exclusively in the interaction term were rare in both species, accounting for only 54 genes in *F. crotonensis* and 16 in *M. aeruginosa* (0.6% and 0.5% of responsive genes, respectively). In *Synechococcus* sp., the two testable effects were largely distinct, with 78.3% of the differentially expressed genes associated exclusively with P limitation (Fig. S3a).

Expression filtering (i.e., excluding genes with low number of reads) retained between 41% (*N. palea*) and 98% (*Synechococcus* sp.) of the genes within each reference genome, for differential expression analysis (Table S1). Functional annotation coverage also varied considerably among species, with only 32.5% of differentially expressed genes in *N. palea* associated with at least one Gene Ontology (GO) term compared with 58.7% in *M. aeruginosa* (Fig. 3a). Detailed differential expression results, including volcano plots, numbers of up- and down-regulated genes, and overlap among contrasts, are provided in Figs. S2 and S3.

### A shared metabolic core in response to warming

To identify common responses to warming across species, we analysed pairwise temperature contrasts (20 vs 24, 24 vs 28 and 20 vs 28 °C) at moderate P limitation (P-mod) and summarised the resulting functional responses across taxa (Fig. 4). *Fragilaria crotonensis* contributes only to the 20 vs 24 °C contrast (Fig. 1c). Results for individual contrasts are shown in Fig. S4. Warming consistently affected broad metabolic functions in all taxa (Fig. 4). Notably, the median proportion of up-regulated genes per category was 47% (Fig. S5).

**Figure 4.**
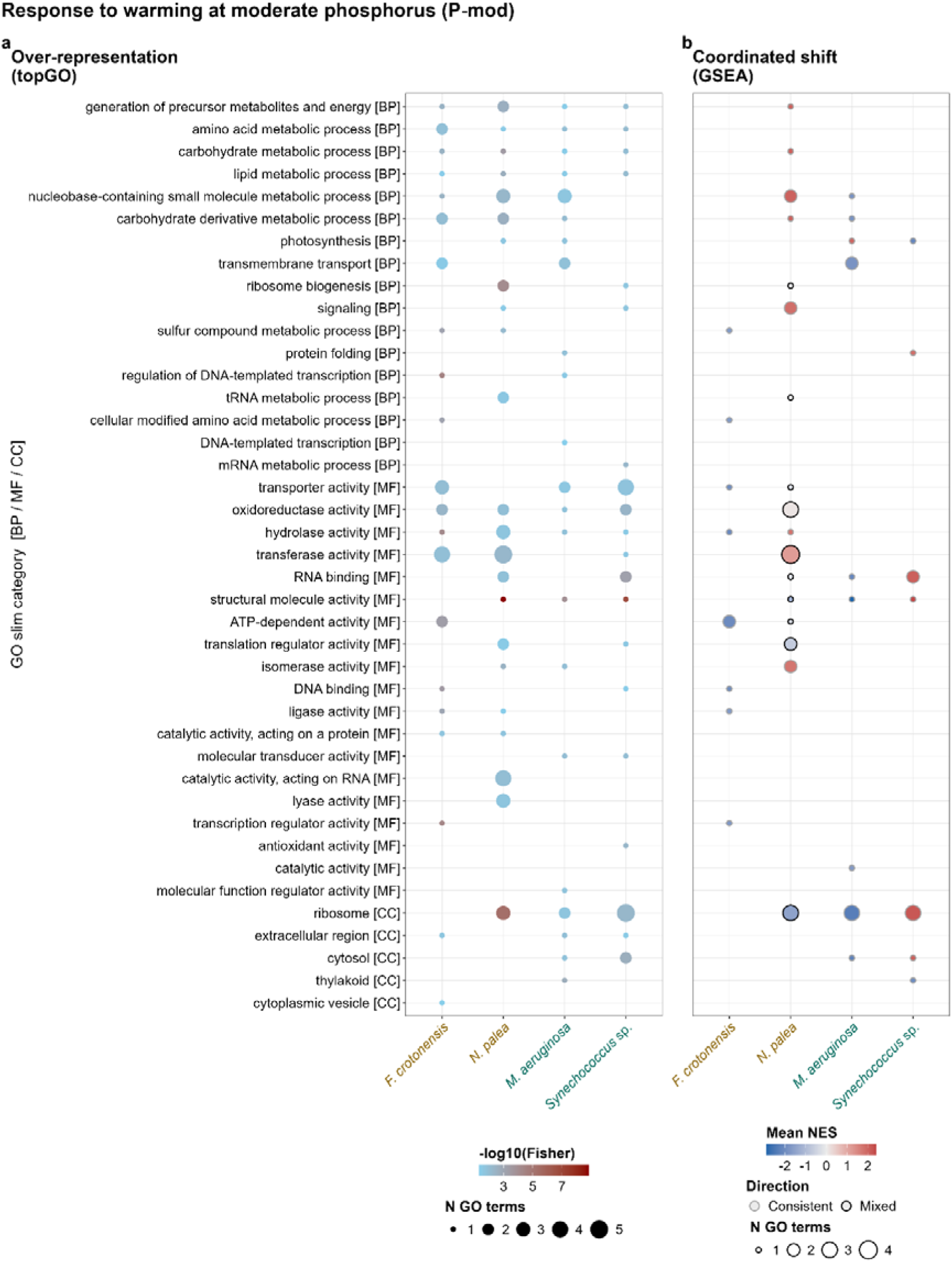
Functional response to warming at moderate phosphorus (P-mod). Results are pooled across pairwise temperature contrasts within each species; *F. crotonensis* contributes only the 24 versus 20 °C comparison. Rows represent Gene Ontology (GO)-slim categories grouped by ontology ([BP] biological process, [MF] molecular function, [CC] cellular component). (a) GO terms over-represented among differentially expressed genes (topGO, Fisher’s exact test, *P* < 0.05). Dot colour indicates mean −log₁₀(*P*) and dot size the number of significant terms. (b) Gene set enrichment analysis (GSEA; adjusted *P* < 0.05). Dot fill indicates mean normalised enrichment score (NES), with positive values indicating higher expression at the warmer temperature. Dot size indicates the number of significant GO terms. Black outlines denote categories containing terms enriched in opposite directions. DEGs were identified at FDR < 0.05 and |log₂ fold change| ≥ 0.5.

### Ribosomal genes exhibited the strongest but contrasting responses to warming

Among all functional categories, ribosome- and translation-related processes showed the strongest and most consistent response to warming (Fig. 4b), remaining significant after FDR correction in three of the four species (Fig. S6a). However, the direction of change differed between species. In *M. aeruginosa*, warming induced a pronounced down-regulation of ribosome-associated functions, generating the largest effect size observed in the study (structural molecule activity, NES = −2.89; ribosome, NES = −2.32; Fig. 4b). These responses were already evident at the 20 vs 24 °C contrast. None of the 22 differentially expressed genes assigned to structural molecule activity increased in expression in this comparison (Fig. S5). In contrast, *Synechococcus* sp. showed the opposite pattern, with coordinated up-regulation of the same categories (Fig. 4b) and all these 25 differentially expressed genes increased in expression at 20 vs 28 °C (Fig. S5). *Nitzschia palea* showed a reversal, with all 16 differentially expressed ribosome biogenesis genes increasing in the 20 vs 24 °C contrast, whereas none of the 14 did in the 24 vs 28 °C contrast (Fig. 4b, S4 and S5). Structural molecule activity in this species gave the strongest enrichment in the study (−log10 FDR = 13.63; Fig. S6a). By contrast, *F. crotonensis* showed no detectable translation-related response. Instead, the warming response was characterized by broad down-regulation of transcriptional and catabolic functions (Fig. 4b), that remained significant after FDR correction (Fig. S6a).

The two cyanobacteria were also distinguished by their photosynthetic response to warming. Photosynthesis was up-regulated in *M. aeruginosa* but down-regulated in *Synechococcus* sp., together with thylakoid-associated functions (Fig. 4b). *Synechococcus* sp. additionally up-regulated protein-folding pathways and, in the 20 vs 28 °C contrast, enriched peroxidase activity and reactive oxygen species metabolic processes (Tables S3, S4). In *N. palea*, signalling and glycolytic, carbohydrate and energy categories were up-regulated (Fig. 4b).

### Phosphorus limitation resulted in divergent responses among species

P limitation generated markedly different transcriptional responses among species (Fig. 5). Positive NES value indicates higher expression under P limitation than under moderate P conditions. The clearest contrast was between the two cyanobacteria, again centred on photosynthesis-related functions (Fig. 5b). In *M. aeruginosa*, P limitation was associated with increased expression of photosynthesis and thylakoid-associated categories (NES = +2.40 and +2.01, respectively), together with energy-generation pathways, but reduced expression of transport, biosynthesis and cell-envelope functions (Fig. 5b). In *Synechococcus* sp., the same treatment produced the opposite pattern, with repression of photosynthesis and thylakoid-related pathways (NES = −2.58 and −2.12, respectively), and enhanced expression of translation- and protein-folding functions. In *F. crotonensis*, P limitation induced a broadly suppressive response (Fig. 5b). All significantly enriched functional categories showed reduced expression, including photosynthesis, transcriptional regulation and transport-related processes, except hydrolase activity with a mixed response (mean NES +0.06). Over-representation analyses showed similar patterns (Fig. 5a). Three enrichments remained significant after FDR correction: transcription regulation in *F. crotonensis*, photosynthesis in *Synechococcus* sp., and thylakoid-associated categories in both cyanobacteria (Fig. S6b). Most P-responsive categories in *F. crotonensis* were detected in the 20 °C comparison but not at 24 °C (in GSEA). In *M. aeruginosa* the response depended on temperature, with transmembrane transport recovered primarily in the 28 °C contrast, but also at 20 °C in GSEA(Fig. S7).

**Figure 5.**
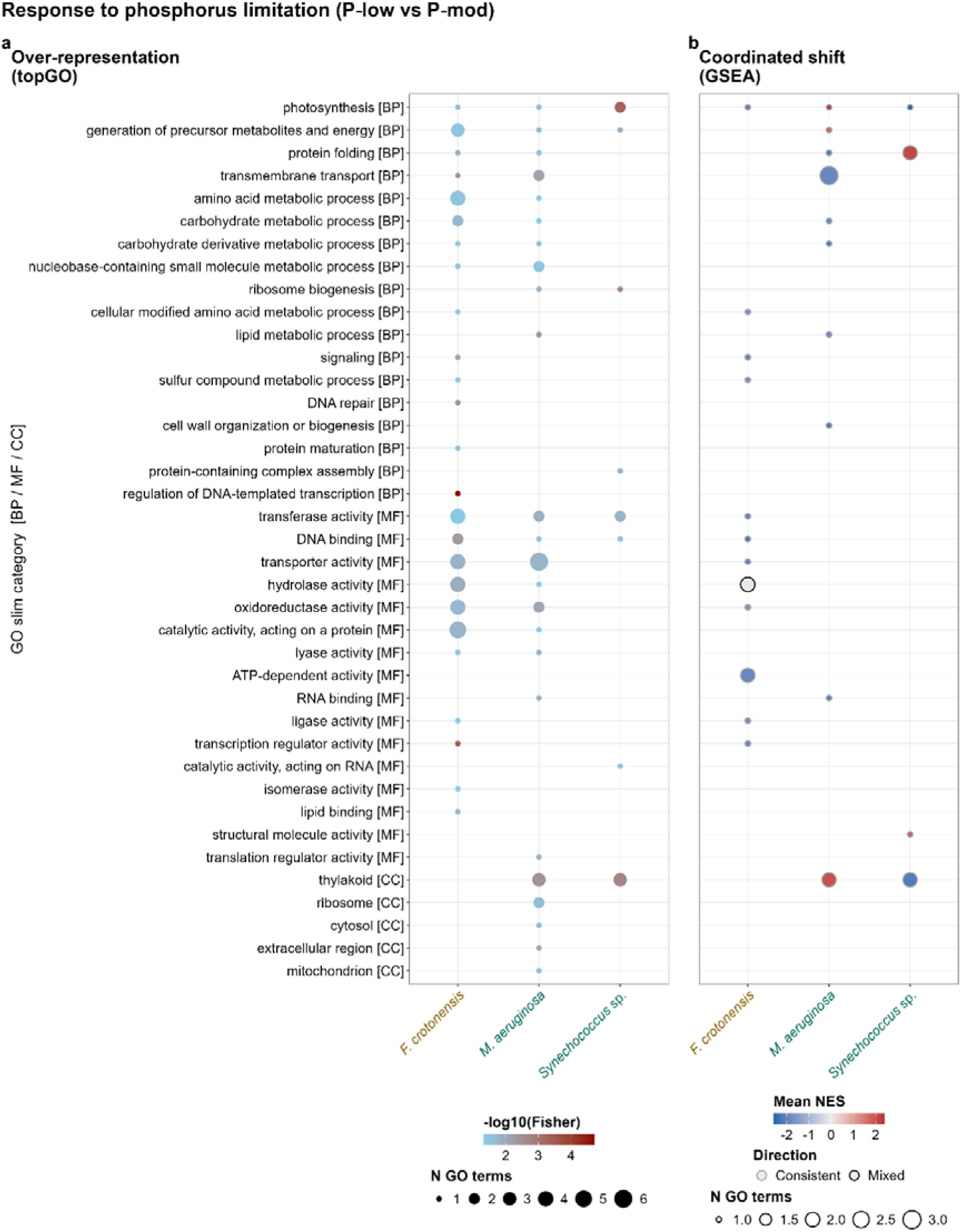
Functional responses to phosphorus limitation. Results are pooled across all temperatures available for each species (*F. crotonensis*: 20 and 24 °C; *M. aeruginosa*: 20, 24, and 28 °C; *Synechococcus* sp.: 24 °C). *N. palea* is omitted because insufficient P-low replicates survived. (a) Gene Ontology (GO) over-representation analysis (topGO) and (b) gene set enrichment analysis (GSEA), displayed as in Figure 4. Positive NES values indicate higher expression under P limitation.

### Phosphorus-acquisition metabolic pathways were not detected

Examination of individual GO terms revealed no enrichment of pathways typically associated with P acquisition, including phosphate transport, phosphatase activity or P-sparing lipid synthesis, in any species (Tables S3, S4). In *M. aeruginosa*, reductions in transport-related functions instead reflected nitrate transport and sulfate transport (Table S4). It should be noted that the annotation coverage of P-related functions was extremely limited across all reference genomes. Only a small portion of expressed genes were assigned to these categories, while categories of cell-cycle regulation and ribosomal function had a greater representation (Table S2). Oxidative stress-related functions responded to P limitation in both *M. aeruginosa* and *F. crotonensis*, including response to oxidative stress pathways (Tables S3, S4).

### Phosphorus supply reshaped warming responses

For the two species where Temperature × Phosphorus interaction could be evaluated, *F. crotonensis* and *M. aeruginosa* (Figs. 1c, 6), interaction effects involved a substantial fraction of expressed genes (2,968 and 1,871 genes, 24.2% and 51.1% of expressed genes, respectively; Fig. 3a), yet relatively few genes responded exclusively to the interaction term (Fig. 3b,c). In *F. crotonensis* functional enrichment analyses identified ATP-dependent activity, transport, transcriptional regulation and hydrolase activity as the categories most strongly affected by the interaction (Fig. 6a), mirroring the dominant responses to warming and P limitation observed separately (Fig. 4b, Fig. 5b). All enriched categories showed negative NES values (−1.88 to −2.14; Fig. 6b), indicating that P limitation weakened the warming response. ATP-dependent activity and hydrolase activity remained significant after FDR correction (Fig. S6c). In *M. aeruginosa*, although no functional category remained significant after FDR correction, GSEA revealed coordinated shifts in ribosomal, energy-generation, carbohydrate-metabolism and cell-wall associated functions, together with positive responses in thylakoid and protein-folding categories (Fig. 6b; Fig. S8). As in *F. crotonensis*, these categories largely correspond to those responding to the main effects of temperature and P (Fig. 4b, Fig. 5b). Antioxidant activity and glutathione biosynthesis were also over-represented among interaction genes for *M. aeruginosa* and *F. crotonensis*, respectively (Fig. 6a).

**Figure 6.**
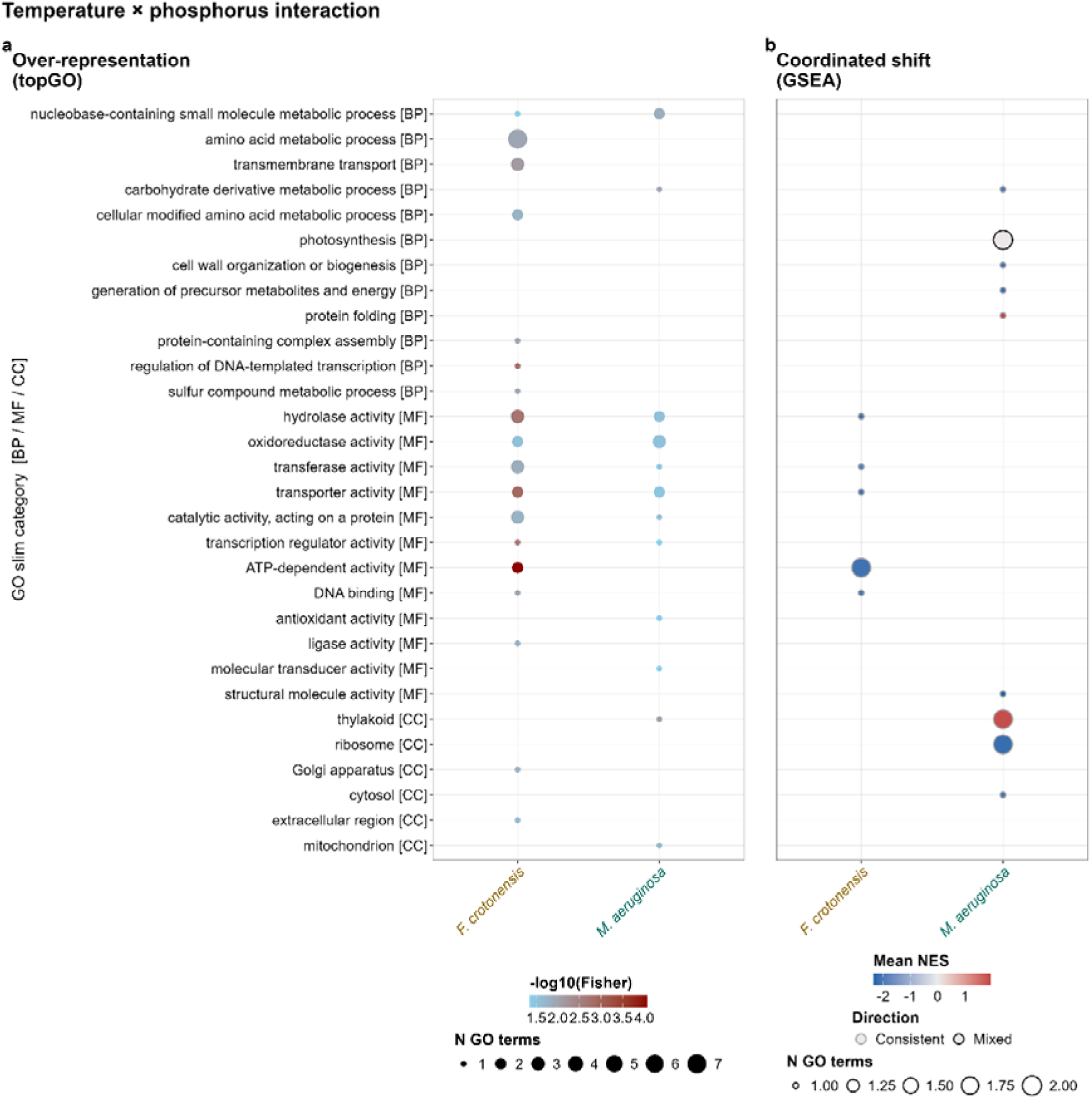
Functional categories affected by the temperature × phosphorus interaction. Results are shown for the two species with full-factorial designs, *F. crotonensis* and *M. aeruginosa*. (a) Gene Ontology (GO) terms over-represented among genes showing a significant interaction in the model-level test (FDR < 0.05). (b) gene set enrichment analysis (GSEA) based on individual interaction coefficients and summarised at the species level. Positive NES values indicate that warming responses were more positive under P-mod than under P limitation. Symbols are as in Figure 4.

Comparing equivalent warming contrasts across P treatments, we observed distinct interaction outcomes between species (Fig. 7). In *F. crotonensis*, P limitation weakened the transcriptional response to warming. Fewer genes responded to the 20 vs 24 °C temperature increase under P limitation than under moderate P conditions (2,378 versus 3,109 genes; Fig. 7c). Of the eight functional categories significantly affected by warming at P-mod, only hydrolase activity remained significant under P limitation, where it changed in the opposite direction (Fig. 7b). In contrast, P limitation amplified warming responses in *M. aeruginosa*. Approximately twice as many genes responded to warming under P limitation as under P-mod conditions, both for the 20 vs 24 °C (1,438 versus 780 genes) and the 20 vs 28 °C comparison (1,482 versus 721 genes; Fig. 7c). The functional nature of the response also changed. Whereas warming responses under moderate P limitation were dominated by ribosome- and translation-related functions (Fig. 4b), responses under P limitation were characterized by changes in energy generation, carbohydrate metabolism, thylakoid organization and protein folding (Fig. 7a, b). Ribosome-related categories instead shifted to positive values in the 20 vs 28 °C contrast under P limitation, the opposite of their response at moderate P, although this rests on two replicates and is only marginally significant (adjusted *p* = 0.04). Photosynthesis-related functions showed both positive and negative shifts depending on the specific GO terms involved (Fig. S9).

**Figure 7.**
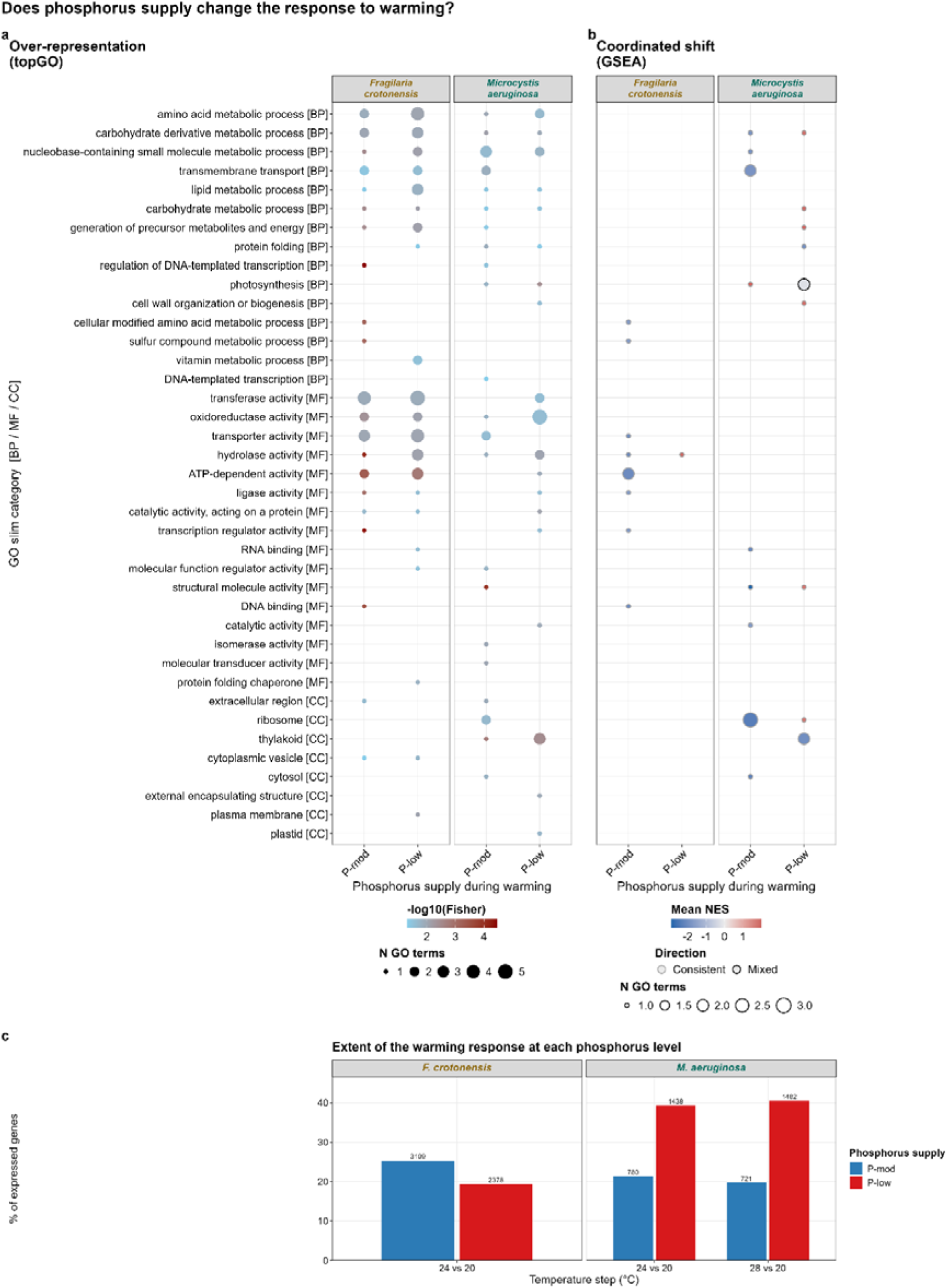
Phosphorus availability reshapes transcriptomic responses to warming. Temperature responses were compared between P-mod and P-low conditions using equivalent temperature contrasts relative to 20 °C (24 vs 20 °C in both species; also 28 vs 20 °C in *M. aeruginosa*) were compared at P-mod and at P-low, with steps pooled within each P level. (a) Gene Ontology (GO) over-representation analysis (topGO) and (b) gene set enrichment analysis (GSEA), displayed as in Figure 4. (c) Differentially expressed genes (DEG) for each temperature step at each P level, as a percentage of expressed genes; labels give the number of DEG. DEG were called at FDR < 0.05 and |log₂ fold change| ≥ 0.5.

## Discussion

Warming and phosphorus (P) limitation engaged many of the same cellular functions across species, but their regulation frequently differed, even within phyla. The divergence therefore lay not in gene content, but in how shared cellular machinery was regulated under environmental stress. Because gene expression is a trait, these differences can be viewed as evolved reaction norms, linking a physiological question – how phytoplankton allocate resources under warming and P limitation – with an evolutionary one – why species of the same phylum regulate the same machinery differently and how these differences may shape future responses to environmental change.

### Ribosomal allocation supports both hypotheses, in different species and temperature ranges

Ecological stoichiometry and thermal physiology make opposite predictions about how warming affects ribosomal investment. The GRH predicts that ribosomal investment rises wherever warming accelerates growth (Elser et al., 2000), whereas the TDPH predicts that it falls because warmer ribosomes work more efficiently (Armin & Inomura, 2021; Toseland et al., 2013). Ribosomal functions gave the strongest response, but its direction varied among species and, in *N. palea*, between temperatures, so each hypothesis was supported only in particular species and temperature ranges.

The response of *N. palea* is the most informative, as the same categories and genes show opposite patterns. Specifically ribosomal and translation-related genes are up-regulated from 20 to 24 °C and down regulated from 24 to 28 °C. Notably, this shift did not occur at the thermal optimum (32.2 °C (95% CI 31.3–33.3 °C under moderate P limitation, 10 µmol L⁻¹;(Levasseur et al., 2025)), but both temperature intervals were on the ascending phase of the thermal performance curve, approximately 4-12 °C below the optimum. Ribosomal investment increased at lower temperatures, consistent with the GRH, but declined at higher temperatures despite growth likely continuing to increase, consistent with the TDPH. One possibility is that the temperature-dependent gains in translational efficiency accumulate with warming and eventually outweigh the increased ribosomal demand associated with faster growth, well before maximum growth is reached. A similar, although not identical, pattern was reported for the bloom-forming dinoflagellate *Prorocentrum shikokuense*, where growth rate increased from 19 to 25 °C and ribosomal protein abundance was higher near the thermal optimum (26.8 °C), whereas warmer conditions induced protein-homeostasis responses and reduced photosystem II efficiency (Zhang et al., 2022). However, unlike *P. shikokuense*, where the decline in ribosomal investment coincided with the thermal optimum, *N. palea* exhibited this shift several degrees below it.

The cyanobacteria only partly conformed to this pattern. *Synechococcus sp.* showed the weakest transcriptional response to warming of the four species. The highest temperature treatment was only slightly above the estimated thermal optimum of *Synechococcus* sp. (Levasseur et al., 2025). Despite this, ribosomal investment increased across the entire temperature range. Because growth just above the thermal optimum is likely still higher than at 20 °C, this pattern is broadly consistent with the GRH. However, ribosomal expression also increased between 24 and 28 °C, a temperature interval likely spanning its thermal optimum, although only five genes contributed to this response. This increase coincided with the up-regulation of protein folding, peroxidase activity and reactive oxygen species metabolism, suggesting that ribosomal expression near and above the optimum may partly reflect protein repair and turnover under heat stress rather than growth. In contrast, *M. aeruginosa* supports the TDPH, since the expression of the same categories was reduced from the first temperature step onward and with the largest effect sizes in this study. For *M. aeruginosa*, growth continued to increase across the temperature range examined, indicating that its thermal optimum likely exceeded the temperatures tested (Levasseur et al., 2025). Its ribosomal decline thus began far below its optimum, as in *N. palea*. *Fragilaria crotonensis* supports neither prediction, showing no translation-related response in any comparison despite carrying the largest repertoire of ribosome- and translation-associated genes among the four species. At P-mod, its thermal optimum is 23.9 °C (95% CI 22.5–25.8 °C) and its upper thermal limit 27.1 °C (Levasseur et al., 2025). The only temperature contrast available for this species (20 vs 24 °C) ends at its thermal optimum, while warmer treatments could not be evaluated because 28 °C exceeds the temperature at which the species can maintain growth. Under these conditions, there may have been little need to adjust ribosomal allocation in either direction. Taken together, these results indicate that thermal position alone cannot predict ribosomal allocation. Instead, species differ in the temperature range over which translational-efficiency gains outweigh growth-associated ribosomal demand: between 24 and 28 °C in *N. palea*, before 24 °C in *M. aeruginosa*, and possibly beyond the temperatures tested here in *Synechococcus* sp.

The contradiction across species with distinct thermal curves may explain why comparable studies disagree. Transcriptomic work on warming in diatoms reports changes in ribosome synthesis and translation without a consistent direction across species (Hong et al., 2025; Thangaraj & Sun, 2021). In addition, a multi-omics study of the freshwater diatom *Cyclotella meneghiniana* under altered nitrogen to P supply likewise found enrichment dominated by the same conserved processes we recover, specifically ribosome-related functions, protein processing and energy metabolism (Liu et al., 2026). If lineages differ in their evolutionary thermal histories, studies that apply similar absolute temperatures to them should be expected to disagree. Our results add that the direction is not conserved even between species of the same phylum or associated clearly with particular thermal optima of growth.

### Alternative strategies under phosphorus limitation

Phosphorus limitation divided the species even more sharply than warming. The clearest example is the response of the photosynthetic apparatus, which was strongly up-regulated under P-limitation in *M. aeruginosa* and strongly down-regulated in *Synechococcus sp.*. The thylakoid categories followed the same pattern. Reducing photosynthetic investment under P-limitation follows the reduced growth demand under P stress (Alipanah et al., 2018). The alternative strategy of sustaining carbon fixation while growth is constrained requires carbohydrate and cell-envelope responses, which is the overall response in *M. aeruginosa*. Similarly, ribosomal investment responses divided the two cyanobacteria. In *M. aeruginosa* P-limitation down-regulated ribosomal responses, which follows the GRH and as reported as well for *T. pseudonana* (Dyhrman et al., 2012). In contrast, *Synechococcus sp.* up-regulated ribosomal categories under P limitation, increasing investment in its most P-demanding processes when P is scarce. This could only be sustained if the cells store P internally, such as polyphosphate, or if they maintained a high cellular P content at the cost of slower growth rate. Otherwise, ribosomal-protein mRNA increased but production of ribosomes did not. Once more, a different response was observed in *F. crotonensis*. Every category recorded under P-limitation was down regulated (except hydrolase activity, which was mixed), and these were the same transcriptional and catabolic categories observed for the temperature response, in the same direction. It seems that this species reduces a common set of regulatory and energy consuming functions in response to both stressors. This may be a response at the gene expression level when a species does not have a strong acquisition response.

From an evolutionary perspective, these are alternative strategies to the same resource constraint. Storage capacity, the balance between acquisition and conservation, and the coupling of carbon fixation to growth rate are all traits that vary among phytoplankton groups and are subject to selection. Alternative strategies like these ones can underpin niche differentiation among co-occurring species (Litchman & Klausmeier, 2008).

### Conserved machinery but divergent regulation

Our study highlights that grouping species by phylum does not predict their responses to temperature and P-limitation. Since photosynthesis, ribosomal investment and chaperone function gene groups are present in all four species, the divergence lies in their regulation rather than in gene content, consistent with the view that gene regulation evolves more readily than the genes it controls (Vigneau et al., 2026; Ye et al., 2022).

### Interaction changes existing responses

P limitation changed the transcriptional response to warming in both species where this interaction effect could be tested, mostly by altering the existing response. Very few genes were differentially expressed only at the interaction level (54 and 16 genes, 0.6% and 0.5%). The two species, however, responded in opposite directions. In *F. crotonensis*, P-limitation suppressed the warming response, as most of the response at moderate P concentrations was not detectable under P-limitation treatment, supporting the constrained pattern we proposed. This species also shows signs of compensation, as the P response decreases with increased temperature, confirming our expectation that these mechanisms would not be mutually exclusive, although only supported by GSEA (Armin & Inomura, 2021; Hessen et al., 2017; Toseland et al., 2013). The constrained pattern found in *F. crotonensis* is supported by the elemental data measured in this experimental system and which found a decrease in the stoichiometric response to temperature under limiting resources (Levasseur et al., accepted). In *M. aeruginosa* we observed the reverse scenario, as warming under P-limitation generated a gene expression response that amplified the response observed under moderate P conditions. Which pattern a species shows may follow from which driver dominates its transcriptome overall: *F. crotonensis*, in which P affected more genes than temperature, was constrained; *M. aeruginosa*, in which temperature dominated, was amplified. Two species cannot establish a rule, but this association is testable. Whether P limitation also shifted the thermal optima of these two species, as it did for *N. palea*, cannot be assessed, because no estimates exist at 0.1 µmol L⁻¹ (Levasseur et al., 2025).

### Annotation bias against group-specific genes

Phosphate uptake acclimates through a sequence of stages along the phosphate continuum, with different transporter classes induced at different points (Cáceres et al., 2019), and the Pho regulon is characteristically a response to acute starvation (Harke & Gobler, 2013). We expected P limitation to induce acquisition and recycling responses through the Pho regulon, including high-affinity phosphate transporters and phosphatases, as reported in marine diatoms (Alipanah et al., 2018; Dyhrman et al., 2012), in *Microcystis aeruginosa* (Harke & Gobler, 2013), in a dinoflagellate (Kalinina et al., 2025), and as detectable in whole lake communities *in situ* (Vila-Costa et al., 2013). In our study no phosphate-specific transport or phosphatase term was significantly enriched. The absence seems to largely reflect limited annotation coverage of P-acquisition genes in the available genomes. Across the expanded set of phosphate transport terms, the number of annotated expressed genes was zero in *F. crotonensis*, one in *N. palea*, nine in *M. aeruginosa* and six in *Synechococcus sp.*, so the set could not be tested in the two diatoms and only reached the minimum size (n=5) in the two cyanobacteria, where no phosphate-transport term was enriched. In contrast, ribosomal and photosynthetic functions were well represented and generated strong signals. In non-model organisms, ontology-based enrichment is often weakest for the lineage-specific gene families most likely to differentiate species ecologically and evolutionarily. Studies that successfully recover the Pho regulon typically rely on manual curation, well-annotated model genomes, or targeted analyses of P-acquisition genes (Alipanah et al., 2018; Dyhrman et al., 2012; Harke & Gobler, 2013; Kalinina et al., 2025). Similarly, P-sparing lipid substitution requires distinguishing sulfolipid and betaine-lipid synthesis from general membrane lipid metabolism, whereas only the latter was annotated here. By contrast, cell-cycle functions were well represented across all species but showed no enrichment, providing an informative null despite previous reports of cell-cycle responses to P deficiency (Kalinina et al., 2025).

### Consequences for predicting community responses to warming and phosphorus limitation

A common practice in biological community and ecosystem models is to aggregate species into broad functional groups assuming that species within a group respond similarly (Litchman et al., 2010; Litchman & Klausmeier, 2008). Our findings question that assumption. The two cyanobacteria exposed to identical treatments showed opposite gene expression responses across three major functional levels, while the two diatoms differed in whether translation responded or not to temperature. Consequently, averaging species within a functional group can hide substantial biological change and may even suggest little response when individual species respond strongly. Which cyanobacterium dominates influences bloom formation, toxin production and food quality for higher trophic levels (Elser et al., 2000; Huisman et al., 2018).

Our results also help explain why molecular divergence is often not reflected at higher levels of biological organization. In this experimental system, P limitation did not alter the temperature sensitivity of population growth (Weber De Melo, Suter, et al., 2025), and stable elemental ratios masked changes in underlying cellular allocation (Levasseur et al., accepted). Yet P strongly modified transcriptional responses to warming. A lack of divergence in higher-level traits should not be interpreted as evidence of similar underlying responses. Because these regulatory differences are heritable, species whose growth or stoichiometric responses look alike may still respond differently to selection under continued environmental change.

In phytoplankton, thermal and nutrient traits appear to combine instead of trade-off, with warm-adapted populations also performing relatively well under resource limitation (Levasseur et al., accepted). Consistent with this pattern, *M. aeruginosa* maintained photosynthetic gene expression under both stressors, whereas *Synechococcus sp.* reduced it under both. Field studies have similarly linked phosphorus acquisition strategies to shifts in cyanobacterial dominance during bloom development (Linz et al., 2025; Lu et al., 2019). Although transcriptional responses alone cannot predict future community composition, they identify the cellular processes through which environmental change may favour some species over others.

### From plastic responses to evolutionary trajectories

Opposite regulation of conserved genes among species of the same phylum is consistent with divergence in how this machinery is regulated, although differences in physiological state, such as each strain’s position relative to its thermal and nutrient optima, could also contribute. The transcriptional shifts observed here represent plastic responses occurring within a generation, but plasticity itself is an evolving trait (Ghalambor et al., 2007). Phytoplankton can adapt rapidly to warming, within about 100 generations, often through evolutionary changes in metabolic and photosynthetic pathways (Padfield et al., 2016; Schaum et al., 2017), while nutrient limitation can lead to evolution of resource requirements (Bernhardt et al., 2020; Gallego & Narwani, 2022) and can limit adaptation to warming (Aranguren-Gassis et al., 2019). In the strains studied here, the thermal optimum at our moderate P limitation (10 µmol L⁻¹) was 2.0 °C lower than at the highest supply (50 µmol L⁻¹) in *Synechococcus* sp., but 4.5 °C higher in *N. palea* and 2.4 °C higher in *F. crotonensis* (Levasseur et al., 2025). Across the full P gradient tested in that study, the optimum of *Synechococcus* sp. declined as P supply decreased, that of *F. crotonensis* increased, and that of *N. palea* remained between about 28 and 32 °C down to 0.5 µmol L⁻¹ before falling to 16.2 °C at 0.1 µmol L⁻¹ (95% CI 1.6–25.0 °C; Levasseur et al. 2025, Fig. S16). Although this last estimate is uncertain, it would place all three of our temperatures above the optimum of P-limited *N. palea*, which may explain why almost no P-low cultures of this species could be sequenced. The same temperature can therefore sit on different parts of a species’ performance curve depending on resource supply.

Species that regulate cellular processes in different ways are therefore likely to have followed different evolutionary trajectories under the same environmental change. Plastic responses may buffer populations from environmental stress, but they may also increase selection if they move phenotypes away from local optima (Chevin et al., 2010). The redistribution of existing responses under combined stress suggests that plastic and evolutionary responses will act largely on the same regulatory machinery. Determining whether the divergence observed here also exists within species is a key next step, as standing intraspecific variation ultimately determines evolutionary potential (Barrett & Schluter, 2008).

### Limitations and future directions

Because we measured transcript abundances, reduced ribosomal-protein transcript levels do not establish reduced ribosome content, rRNA abundance or cellular P demand, and the predictions of Toseland et al. (Toseland et al., 2013) and Armin and Inomura (Armin & Inomura, 2021) concern macromolecular allocation rather than gene expression. In the freshwater diatom *Cyclotella meneghiniana*, transcript and protein responses to altered N:P supply were only partly coupled (Liu et al., 2026). At the level of elemental composition, C:P and N:P ratios increased with temperature under replete P conditions across the nine species of the companion experiment (Levasseur et al., accepted), consistent with the TDPH. Our results are consistent with these patterns but do not test them directly.

Two further limitations apply. First, each species was represented by a single strain, so the divergence we report is between strains and not demonstrably between species. Second, sparse annotation of lineage-specific gene families left the predicted P-acquisition responses largely untestable.

Four steps would turn these observations into testable predictions. First, warming responses should be interpreted relative to each species’ thermal performance curve, measured under the same resource conditions, with temperatures on both sides of the optimum. The *N. palea* reversal shows that a single warming step, within the ascending phase of the thermal optimum, can give either answer. Second, functional groups should be treated as a hypothesis to test, by quantifying within-group variance before aggregating species and by testing drivers in factorial combination. Third, multiple strains per species, experimental evolution under combined drivers and direct measurements of cellular RNA and elemental composition would show whether the regulatory divergence we report is fixed between lineages and how it translates into cellular allocation. Fourth, targeted analysis of the Pho regulon and other P acquisition and recycling genes, through curated annotation, qPCR or proteomics, would test the predictions that annotation left untestable here.

## Conclusion

Species can possess similar molecular machinery while using it in fundamentally different ways. *A*cross all four phytoplankton species, molecular responses to warming and P limitation were highly lineage-specific, even among taxa belonging to the same functional group. Conserved cellular processes were frequently regulated in opposite directions under identical environmental conditions, and interactions between warming and P limitation primarily reshaped existing transcriptional programmes rather than recruiting novel functional pathways. Ribosomal allocation, the trait most directly linking temperature to cellular P demand, responded neither uniformly nor monotonically. The GRH, which predicts more ribosomes to sustain faster growth, and the TDPH, which predicts fewer, because warmer ribosomes work more efficiently, were both supported, sometimes within the same species. In *N. palea*, the switch between them occurred well below the thermal optimum. Thermal position alone could not explain the patterns observed in the other species, and thermal optima themselves varied with P availability. Because thermal optima are both plastic and evolutionarily labile, and because selection can act directly on regulatory mechanisms, forecasts of phytoplankton responses to global change are likely to be more robust when based on lineage-specific thermal and nutrient traits measured under realistic combinations of environmental stressors, rather than on functional-group averages alone.

## Acknowledgements

We thank Marta Reyes and Oliver Hansen for the lab sampling, Marco Giulio and Max Hofland for conducting the molecular laboratory work, and Niklaus Zemp for bioinformatics support. Data produced and analysed in this paper were generated in collaboration with the Genetic Diversity Centre (GDC), ETH Zurich.

## Data Accessibility

Raw RNA-seq reads, processed count tables, gene-to-GO annotation tables and all analysis scripts are deposited in the ERIC Open Research Data repository (DOI: xxx).

## Benefit-Sharing

Benefits from this research accrue from the sharing of our data and results on public databases as described above.

## Author Contributions

JLS: Data curation, Formal analysis, Software, Project administrations, Visualization, Writing- original draft, Writing- review and editing. VWDM: Conceptualization, Investigation, data curation, Formal analysis, Project administration, Writing – original draft, Writing – review and editing. SL: Investigation, Writing – review and editing. AATW: Methodology, Supervision, Funding acquisition, Writing – review and editing. AN: Conceptualization, Supervision, Funding acquisition, Resources, Writing – original draft, Writing – review and editing.

