## Supplementary Material for "Divergent transcriptional responses of conserved cellular machinery to warming and phosphorus limitation in freshwater phytoplankton"

*Shared first author contribution

+Shared last author contribution

**Appendix S1. Supplementary methods**

**Acclimation and culture conditions**. Prior to each experiment, phytoplankton populations were acclimated to temperature and P conditions in two sequential steps. Temperature acclimation was performed first: the culture temperature was raised by 2 °C per day over four days until the target temperature was reached, while cultures assigned to 20 °C remained constant. Populations were then maintained at the target temperature for a further 48 h. Phosphorus (P) acclimation followed, suring which cultures were split, centrifuged to remove residual nutrients and, resuspended in medium adjusted to the target P concentration. This procedure minimised nutrient carry-over and allowed physiological adjustment to the nutrient limitation imposed.

Experiments were conducted in sterile, ventilated 500 mL tissue-culture flasks, each containing a single species at one combination of temperature and P limitation. Low-biomass inoculation was standardised by pigment fluorescence measured in acclimated cultures on 200 µL subsamples with an Agilent Cytation 5 plate reader, with chlorophyll fluorescence (Ex/Em 445/685 nm) for diatoms and phycocyanin fluorescence (Ex/Em 586/647 nm) for cyanobacteria. Cultures were diluted to a target inoculum of 20 RFU in 350 mL of COMBO medium. P limitation was imposed through medium composition, and cultures were maintained in temperature-controlled incubators under continuous shaking (70 rpm) under an 18L:6D light cycle. Growth was monitored daily by fluorescence until cultures approached steady state, or for a maximum of ten days. All inoculations and sampling procedures were conducted under sterile conditions.

**Normalisation offset**. Effective transcript lengths were divided by their geometric mean across samples, and counts were divided by the resulting scaled length matrix. Effective library sizes were then estimated from the length-scaled counts with the TMM method implemented in calcNormFactors (Robinson et al., 2010; Chen et al., 2025). The length matrix was then multiplied by these effective library sizes, log-transformed and supplied to the DGEList through scaleOffset. Because the TMM normalisation factors are incorporated into the offset, TMM normalisation was not applied a second time.

**Ordinations**. Principal component analysis (PCA) and multidimensional scaling (MDS) included all samples for a species, including those excluded from fitted models, so that variation associated with both temperature and P treatments contributed to the ordination space. Counts were filtered and normalised as for the differential expression analysis, converted to log₂ counts per million using a prior count of 1, and genes with zero variance were removed. PCA was performed with prcomp (base R) on scaled, transposed log-CPM values. MDS was performed with plotMDS (limma), where distances are root-mean-square log₂ fold changes over the 500 most variable genes for each pair of samples.

**GO over-representation: analytical choices.** The weight01 algorithm accounts for dependencies among GO terms, and the minimum node size of five avoids testing very small, uninformative categories. The three GO domains were tested separately, because a topGO object covers one domain at a time, with the same gene-to-GO mapping supplied to all three. As these represent separate families of tests, *p*-values were adjusted within, rather than across domains. Because weight01 algorithm conditions each term on neighbouring terms in the GO graph, the resulting *p*-values are neither independent nor uncorrected in the conventional sense, making additional Benjamini–Hochberg adjustment across terms is conservative. Nominal Fisher *p*-values were therefore used for the main figures, and terms significant after FDR correction are shown in Fig. S6. GO terms with Fisher *p* < 0.05 were used for the GO-slim summaries because few terms remained significant after FDR correction.

**Direction of change within enriched categories**. Over-representation was not tested separately for up- and down-regulated genes because dividing the gene set reduces statistical power. Moreover, transcript-level direction does not necessarily imply the direction of effect on a biological process: genes may function at pathway branch points, catalyse non-limiting steps, or encode negative regulators. To describe, rather than test, the composition of enriched categories, we counted the numbers of differentially expressed genes showing increased and decreased expression and report these proportions in Fig. S5. This descriptive summary involves no additional statistical test and therefore requires no multiple-testing correction.

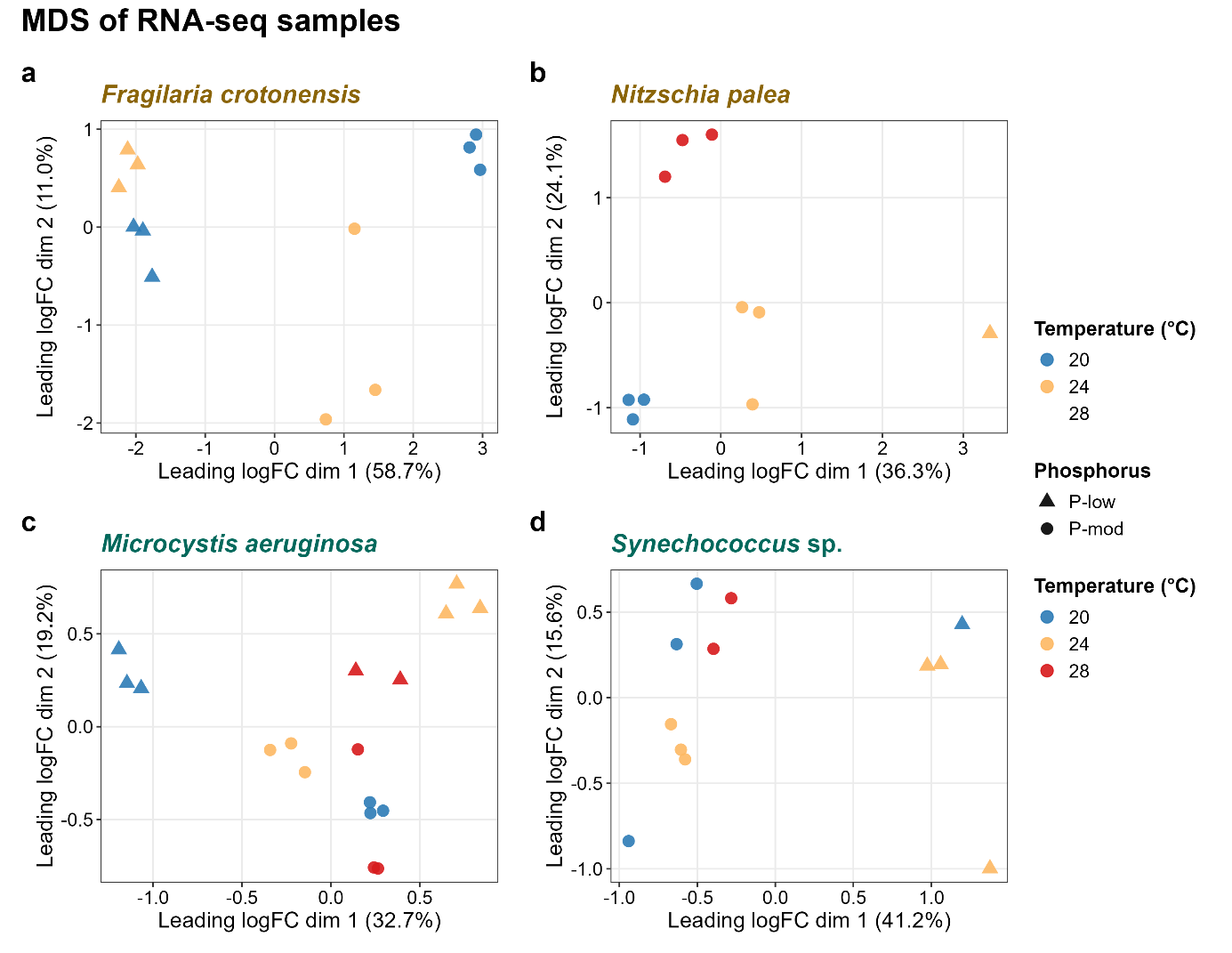
**Figure S1. Multidimensional scaling (MDS) of RNA-seq samples.** MDS ordinations for (a) *F. crotonensis*, (b) *N. palea*, (c) *M. aeruginosa*, and (d) *Synechococcus* sp. Distances represent leading log₂ fold changes calculated from the 500 most variable genes. Colours indicate temperature and symbols indicate P availability (circles, moderate P (P-mod); triangles, P-low). All sequenced samples are shown.

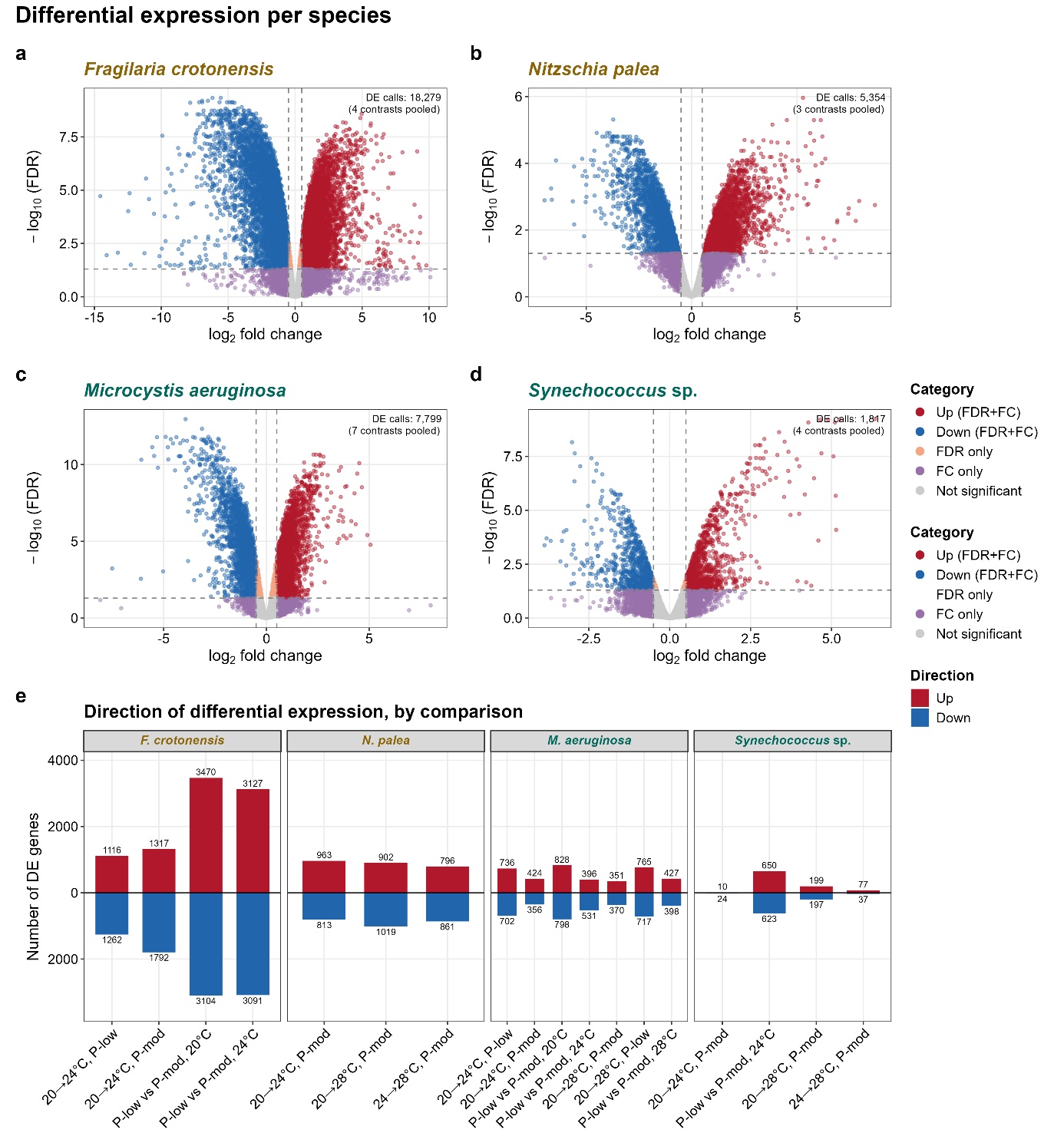
**Figure S2. Differential expression across species and contrasts.** (a-d) Volcano plots for *F. crotonensis*, *N. palea*, *M. aeruginosa*, and *Synechococcus* sp., respectively. Results from all pairwise contrasts are combined; genes significant in multiple contrasts appear once per contrast. Dots are coloured according to significance criteria (FDR < 0.05 and/or |log₂ fold change| ≥ 0.5). Dashed lines indicate significance thresholds. Labels give the number of differentially expressed (DE) genes (a gene DE in several contrasts is counted once per contrast) and of contrasts pooled. (e) Numbers of up- and down-regulated genes in each pairwise contrast. Positive values indicate higher expression in the warmer temperature or under P limitation.

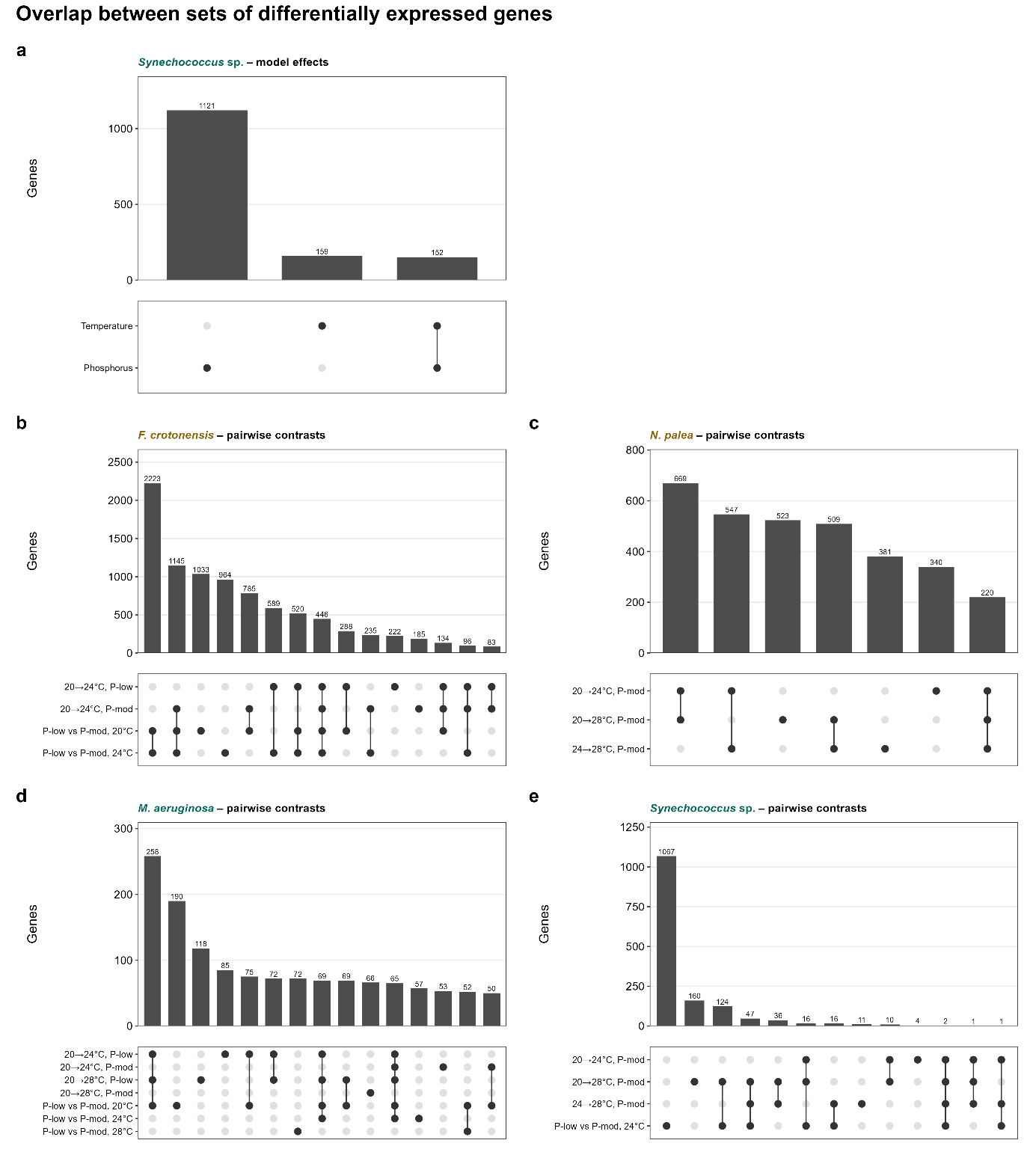
**Figure S3.** **Overlap among differentially expressed genes (DEG)**. (a) Overlap between DEGs associated with temperature and phosphorus (P) availability from model approach in *Synechococcus* sp. (b-e) Overlap among pairwise contrasts within *F. crotonensis*, *N. palea*, *M. aeruginosa*, and *Synechococcus* sp., respectively. Bars indicate the number of genes in each intersection and filled dots indicate the contributing contrasts. Comparisons are restricted to within-species overlaps because gene identifiers are not shared among species.

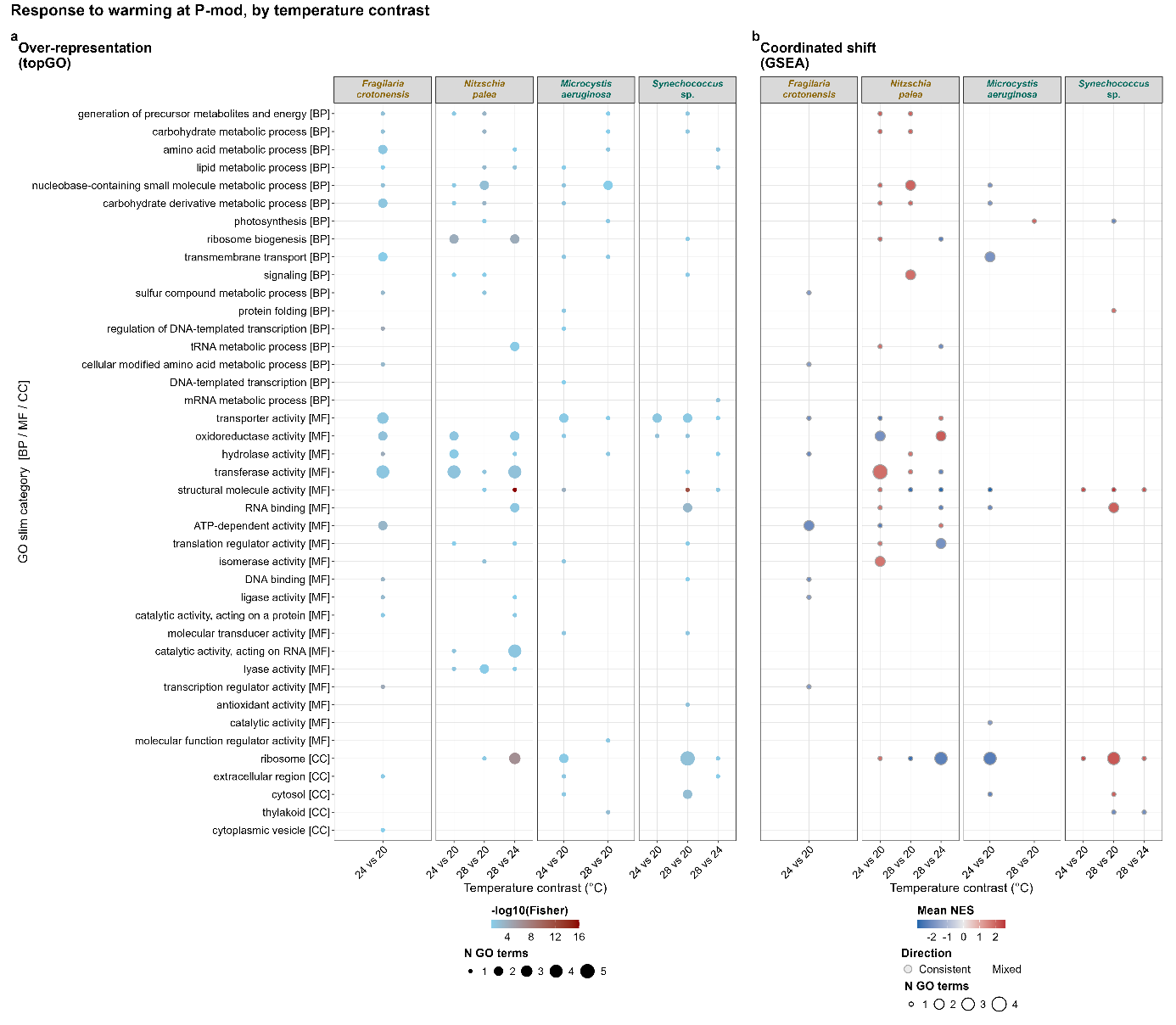
**Figure S4.** **Functional responses to warming under moderate phosphorus availability, by temperature contrast.** Columns represent individual temperature contrasts within each species. (a) Gene Ontology (GO) over-representation analysis (topGO). Dot colour indicates mean −log₁₀(*P*) and dot size the number of significant terms. (b) Gene set enrichment analysis (GSEA). Dot fill indicates mean NES, with positive values indicating higher expression at the warmer temperature. Black outlines denote categories containing terms enriched in opposite directions. Rows represent GO-slim categories.

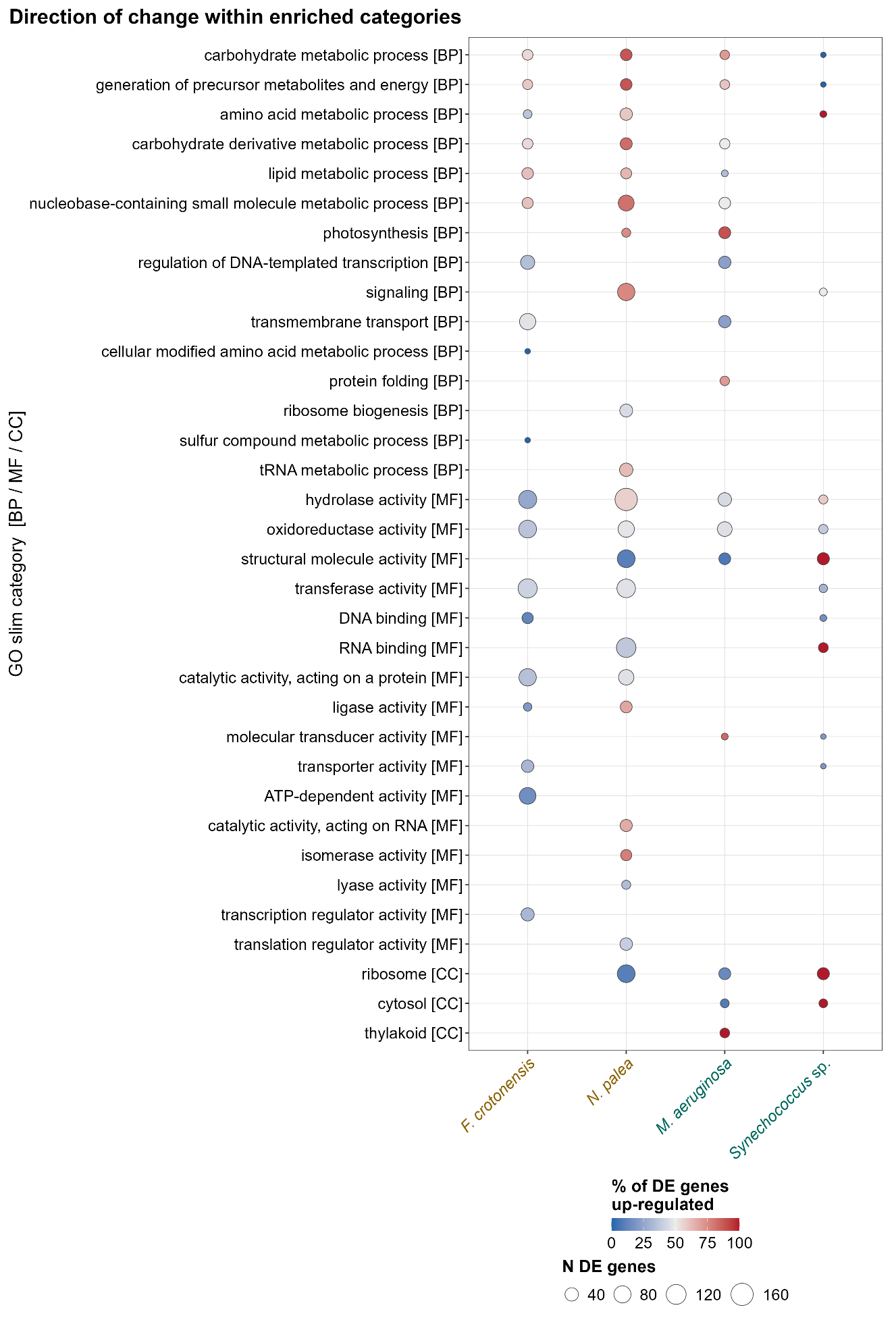
**Figure S5.** **Direction of differential expression within GO-slim categories responding to warming.** Dot fill indicates the percentage of DEGs within a category showing increased expression at the warmer temperature, and dot size indicates the number of DEGs in the category. Categories with fewer than five DEGs are omitted. This figure is descriptive and does not involve formal statistical testing.

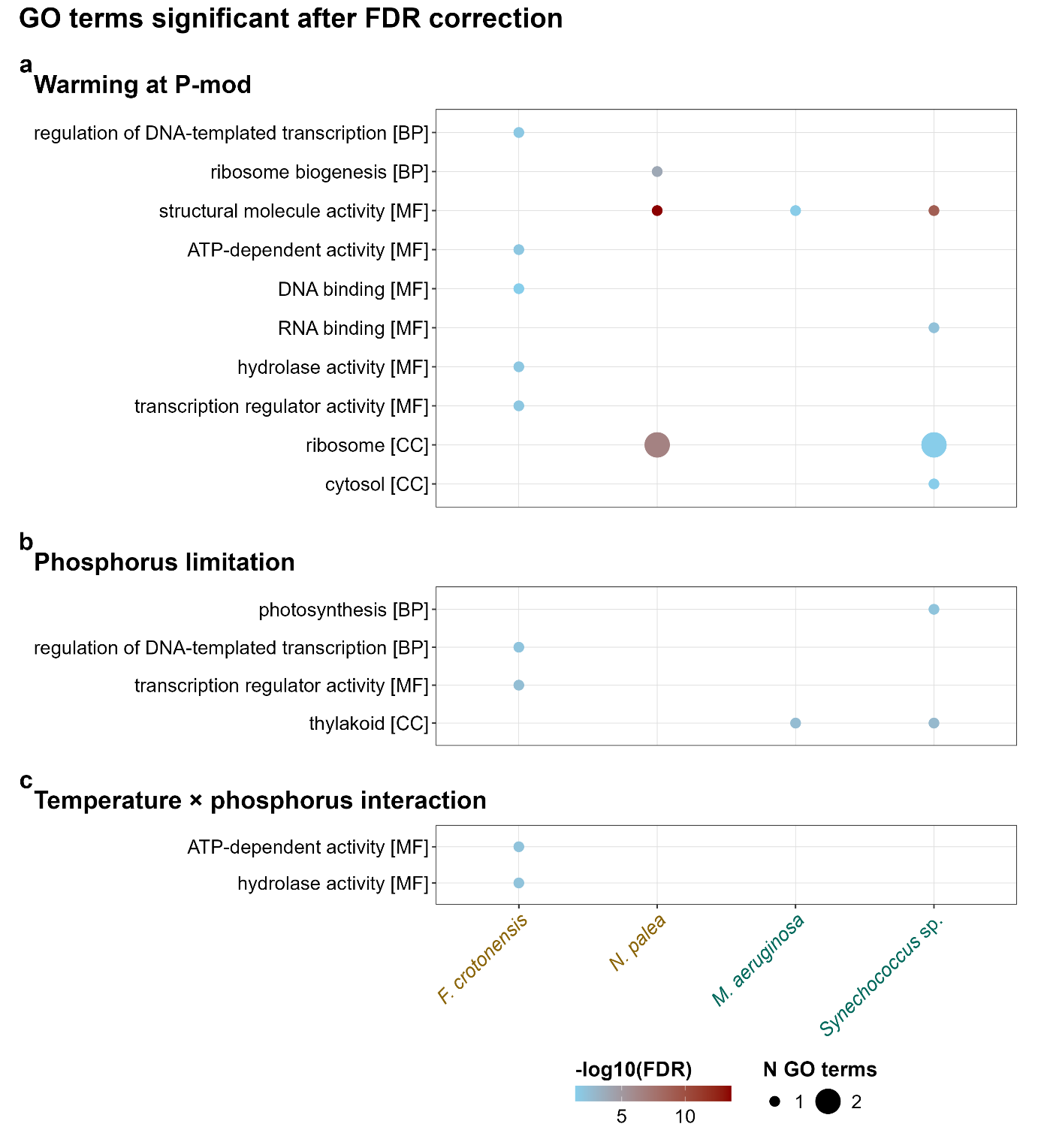
**Figure S6.** **GO-slim categories containing GO terms significant after multiple-testing correction.** Categories include GO terms significant following Benjamini-Hochberg correction (FDR < 0.05) for (a) warming under P-mod, (b) phosphorus limitation, and (c) the temperature × phosphorus interaction. Dot colour indicates mean −log₁₀(FDR) and dot size the number of significant terms.

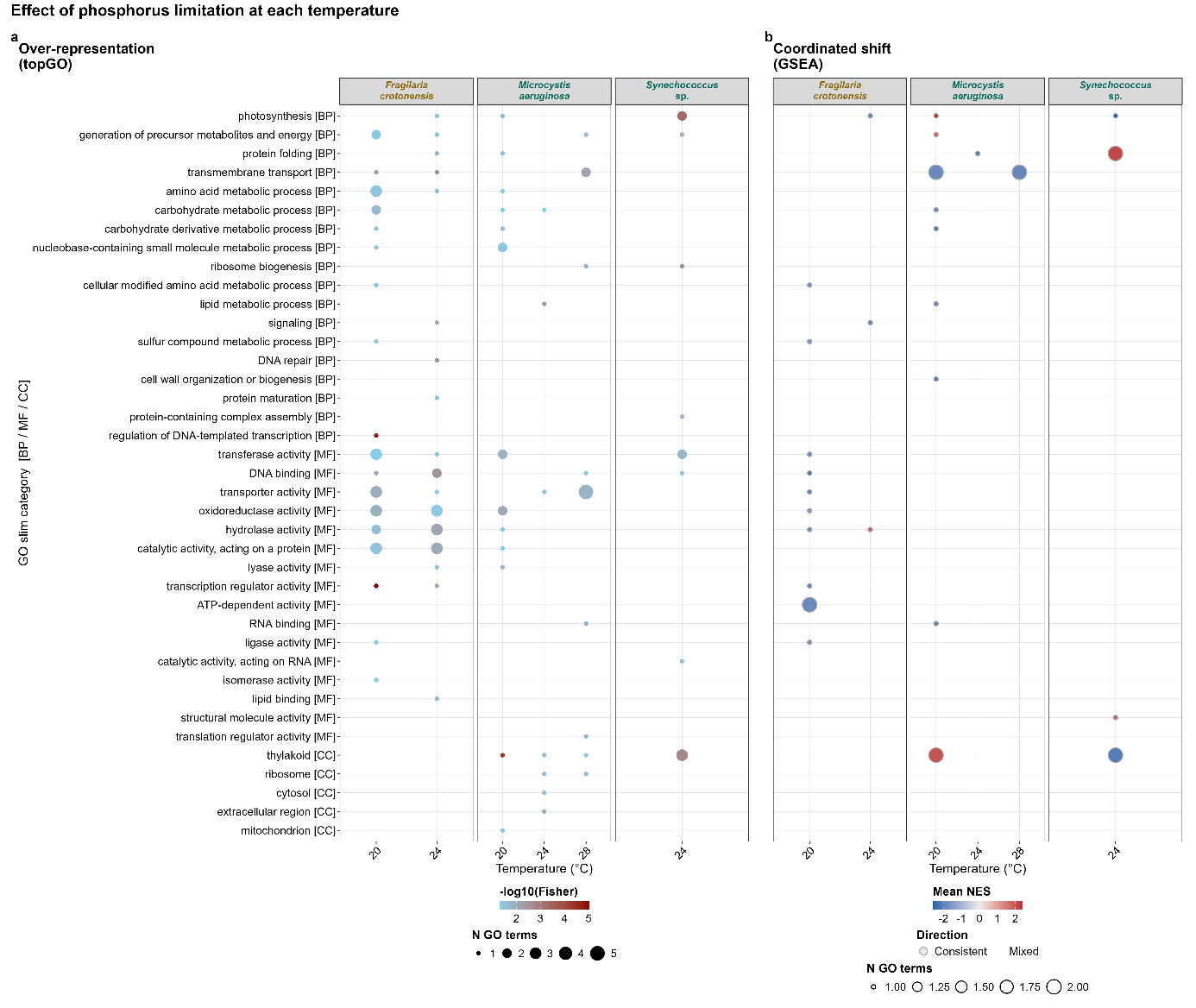
**Figure S7.** **Functional response to phosphorus limitation at each temperature**. Columns represent temperatures at which P-low and P-mod treatments were compared. (a) GO over-representation analysis (topGO) and (b) Gene set enrichment analysis (GSEA), displayed as in Figure S4. Positive NES values indicate higher expression under P limitation.

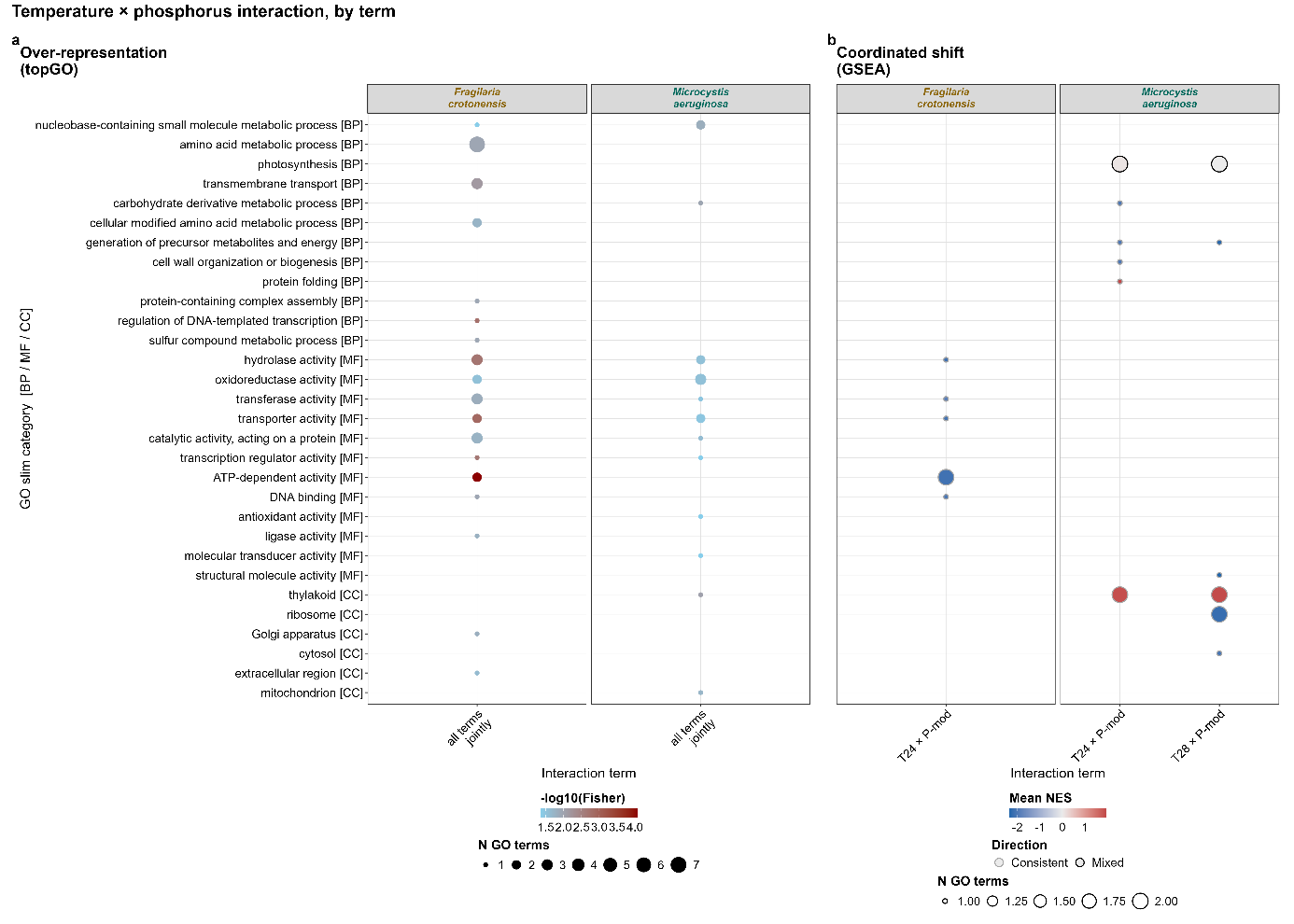
**Figure S8. Functional categories associated with temperature × phosphorus interactions.** (a) GO over-representation analysis (topGO) based on the model-level interaction test. (b) Gene set enrichment analysis (GSEA) performed separately for each interaction coefficient (24 °C × P-mod; and 28 °C × P-mod in *M. aeruginosa*). Displayed as in Figure S4.

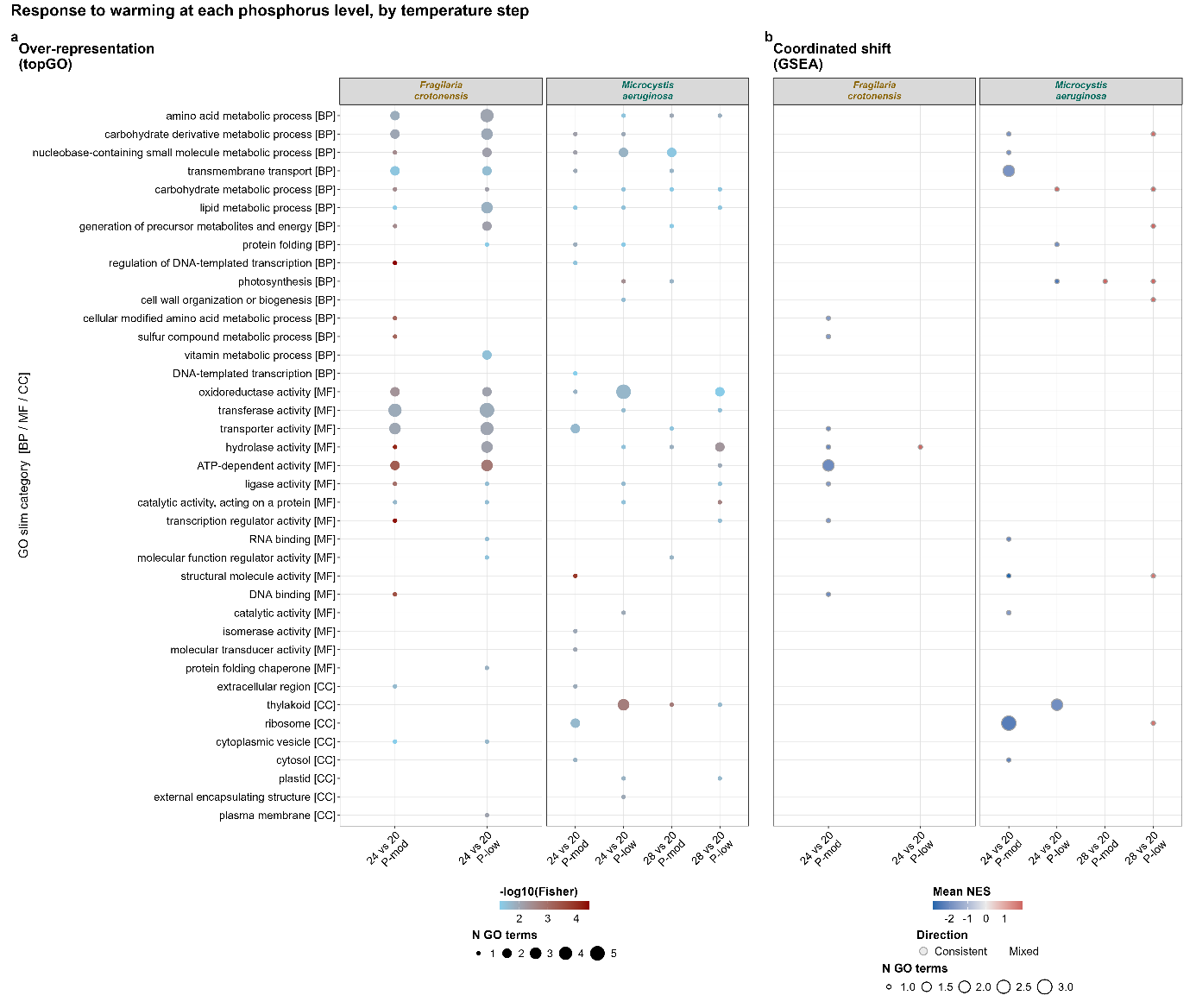
**Figure S9.** **Functional responses to warming at different phosphorus availabilities.** Columns represent combinations of temperature contrast and P limitation (P-mod or P-low). (a) GO over-representation analysis (topGO) and (b) Gene set enrichment analysis (GSEA), displayed as in Figure S4. Positive NES values indicate higher expression at the warmer temperature.

**Table S1. Sequenced samples and gene counts.** Sequenced samples, including species, temperature (T (°C)), P limitation (P-mod, 10 µmol L⁻¹; P-low, 0.1 µmol L⁻¹), number of biological replicates (R) per treatment, Sequencing Lane (Lane A, low-yield libraries; Lane B, high-yield libraries), inclusion in differential expression analysis (≥2 replicates) (DEG); filtered reads pairs in millions (F. read pairs); prcentage of pseudoaligned reads (Align).

| Sample | Sequence ID | Species | T (°C) | P limitation | R | SL | DEG | F. read pairs | Align (%) |
| --- | --- | --- | --- | --- | --- | --- | --- | --- | --- |
| sample46 | 344931_08 | *Fragilaria crotonensis* | 20 | P-mod | 1 | Lane B | yes | 31.39 | 28.4 |
| sample47 | 344931_09 | *Fragilaria crotonensis* | 20 | P-mod | 2 | Lane B | yes | 45.35 | 29.1 |
| sample48 | 344931_10 | *Fragilaria crotonensis* | 20 | P-mod | 3 | Lane B | yes | 37.59 | 28.2 |
| sample100 | 344931_20 | *Fragilaria crotonensis* | 24 | P-mod | 1 | Lane B | yes | 34.01 | 33.4 |
| sample101 | 344931_21 | *Fragilaria crotonensis* | 24 | P-mod | 2 | Lane B | yes | 47 | 32.1 |
| sample102 | 344931_22 | *Fragilaria crotonensis* | 24 | P-mod | 3 | Lane B | yes | 44.15 | 32.5 |
| sample19 | 344932_01 | *Fragilaria crotonensis* | 20 | P-low | 1 | Lane A | yes | 27.91 | 31.2 |
| sample20 | 344932_02 | *Fragilaria crotonensis* | 20 | P-low | 2 | Lane A | yes | 30.4 | 31 |
| sample21 | 344932_03 | *Fragilaria crotonensis* | 20 | P-low | 3 | Lane A | yes | 30.35 | 30.2 |
| sample73 | 344932_14 | *Fragilaria crotonensis* | 24 | P-low | 1 | Lane A | yes | 25.42 | 23.3 |
| sample74 | 344932_15 | *Fragilaria crotonensis* | 24 | P-low | 2 | Lane A | yes | 25.41 | 26.4 |
| sample75 | 344932_16 | *Fragilaria crotonensis* | 24 | P-low | 3 | Lane A | yes | 28.91 | 27 |
| sample43 | 344931_05 | *Nitzschia palea* C | 20 | P-mod | 1 | Lane B | yes | 29.76 | 29.7 |
| sample44 | 344931_06 | *Nitzschia palea* C | 20 | P-mod | 2 | Lane B | yes | 15.65 | 29.7 |
| sample45 | 344931_07 | *Nitzschia palea* C | 20 | P-mod | 3 | Lane B | yes | 26.48 | 30 |
| sample97 | 344932_25 | *Nitzschia palea* C | 24 | P-mod | 1 | Lane A | yes | 24.19 | 33.3 |
| sample98 | 344932_26 | *Nitzschia palea* C | 24 | P-mod | 2 | Lane A | yes | 20.6 | 33.5 |
| sample99 | 344932_27 | *Nitzschia palea* C | 24 | P-mod | 3 | Lane A | yes | 24.28 | 34.3 |
| sample151 | 344932_32 | *Nitzschia palea* C | 28 | P-mod | 1 | Lane A | yes | 30.02 | 30.5 |
| sample152 | 344932_33 | *Nitzschia palea* C | 28 | P-mod | 2 | Lane A | yes | 25.19 | 33.3 |
| sample153 | 344931_27 | *Nitzschia palea* C | 28 | P-mod | 3 | Lane B | yes | 31.46 | 34 |
| sample72 | 344932_13 | *Nitzschia palea* C | 24 | P-low | 3 | Lane A | no | 22.45 | 6.7 |
| sample52 | 344931_13 | *Microcystis aeruginosa* (strain 1) | 20 | P-mod | 1 | Lane B | yes | 17.54 | 45.9 |
| sample53 | 344931_14 | *Microcystis aeruginosa* (strain 1) | 20 | P-mod | 2 | Lane B | yes | 21.86 | 47 |
| sample54 | 344931_15 | *Microcystis aeruginosa* (strain 1) | 20 | P-mod | 3 | Lane B | yes | 19.02 | 48.3 |
| sample106 | 344931_25 | *Microcystis aeruginosa* (strain 1) | 24 | P-mod | 1 | Lane B | yes | 14.04 | 34.9 |
| sample107 | 344931_26 | *Microcystis aeruginosa* (strain 1) | 24 | P-mod | 2 | Lane B | yes | 26.37 | 45.9 |
| sample108 | 344932_29 | *Microcystis aeruginosa* (strain 1) | 24 | P-mod | 3 | Lane A | yes | 16.9 | 42.1 |
| sample160 | 344931_31 | *Microcystis aeruginosa* (strain 1) | 28 | P-mod | 1 | Lane B | yes | 13.82 | 48 |
| sample161 | 344931_32 | *Microcystis aeruginosa* (strain 1) | 28 | P-mod | 2 | Lane B | yes | 14.48 | 51.3 |
| sample162 | 344931_33 | *Microcystis aeruginosa* (strain 1) | 28 | P-mod | 3 | Lane B | yes | 16.49 | 50.3 |
| sample25 | 344932_05 | *Microcystis aeruginosa* (strain 1) | 20 | P-low | 1 | Lane A | yes | 33.36 | 56.9 |
| sample26 | 344932_06 | *Microcystis aeruginosa* (strain 1) | 20 | P-low | 2 | Lane A | yes | 30.64 | 56.4 |
| sample27 | 344932_07 | *Microcystis aeruginosa* (strain 1) | 20 | P-low | 3 | Lane A | yes | 37.68 | 58.1 |
| sample79 | 344932_20 | *Microcystis aeruginosa* (strain 1) | 24 | P-low | 1 | Lane A | yes | 33.14 | 52.5 |
| sample80 | 344932_21 | *Microcystis aeruginosa* (strain 1) | 24 | P-low | 2 | Lane A | yes | 47.61 | 44.1 |
| sample81 | 344932_22 | *Microcystis aeruginosa* (strain 1) | 24 | P-low | 3 | Lane A | yes | 40.28 | 51.5 |
| sample133 | 344932_30 | *Microcystis aeruginosa* (strain 1) | 28 | P-low | 1 | Lane A | yes | 64.73 | 50.2 |
| sample135 | 344932_31 | *Microcystis aeruginosa* (strain 1) | 28 | P-low | 3 | Lane A | yes | 59 | 50.6 |
| sample49 | 344931_11 | *Synechococcus sp.* (CCAP 1478/11) | 20 | P-mod | 1 | Lane B | yes | 35.03 | 47.2 |
| sample50 | 344932_12 | *Synechococcus sp.* (CCAP 1478/11) | 20 | P-mod | 2 | Lane A | yes | 28.01 | 63.8 |
| sample51 | 344931_12 | *Synechococcus sp.* (CCAP 1478/11) | 20 | P-mod | 3 | Lane B | yes | 20.92 | 62.6 |
| sample103 | 344931_23 | *Synechococcus sp.* (CCAP 1478/11) | 24 | P-mod | 1 | Lane B | yes | 15.27 | 27.1 |
| sample104 | 344932_28 | *Synechococcus sp.* (CCAP 1478/11) | 24 | P-mod | 2 | Lane A | yes | 22.98 | 44.4 |
| sample105 | 344931_24 | *Synechococcus sp.* (CCAP 1478/11) | 24 | P-mod | 3 | Lane B | yes | 14.7 | 46.4 |
| sample157 | 344931_28 | *Synechococcus sp.* (CCAP 1478/11) | 28 | P-mod | 1 | Lane B | yes | 14.83 | 43.9 |
| sample158 | 344931_29 | *Synechococcus sp.* (CCAP 1478/11) | 28 | P-mod | 2 | Lane B | yes | 17.34 | 42.9 |
| sample22 | 344932_04 | *Synechococcus sp.* (CCAP 1478/11) | 20 | P-low | 1 | Lane A | no | 44.68 | 27.5 |
| sample76 | 344932_17 | *Synechococcus sp.* (CCAP 1478/11) | 24 | P-low | 1 | Lane A | yes | 26.65 | 2.8 |
| sample77 | 344932_18 | *Synechococcus sp.* (CCAP 1478/11) | 24 | P-low | 2 | Lane A | yes | 20.05 | 72.4 |
| sample78 | 344932_19 | *Synechococcus sp.* (CCAP 1478/11) | 24 | P-low | 3 | Lane A | yes | 27.83 | 72 |

**Table S2. Annotation coverage of the gene sets central to our predictions.** Gene sets included ribosome and translation, phosphate transport, phosphatases, cell cycle and DNA replication, oxidative stress, membrane lipids, and photosynthesis (control). Each gene set comprised seed Gene Ontology (GO) terms and all descendant terms. Values indicate the number of expressed genes annotated to each gene set in each species. Gene sets containing fewer than five genes (the minimum node size) were not eligible for enrichment analyses.

**Table S3. GO terms over-represented among differentially expressed genes.** Significant results from topGO (weight01 algorithm; Fisher’s exact test, *P* < 0.05) for all model effects and pairwise contrasts in each species. For each term: species, effect or contrast, ontology, GO identifier and name, assigned GO slim category, number of annotated genes, observed and expected numbers of differentially expressed genes, Fisher’s *P*-value, Benjamini-Hochberg FDR within each effect and ontology, and whether the term is displayed in the figures. Terms lacking a GO slim ancestor (195 of 808 significant results across Tables S3 and S4) are reported only in the supplementary tables.

**Table S4. GO gene sets significant in gene set enrichment analysis (GSEA).** Significant GO gene sets (5-500 genes; Benjamini-Hochberg FDR < 0.05) from all pairwise contrasts and interaction coefficients in each species. For each gene set: species, contrast, ontology, GO identifier and name, assigned GO slim category, gene set size, normalized enrichment score (NES; positive values indicate higher expression in the tested level), nominal *P*-value, FDR, number of leading-edge genes, and whether the gene set is displayed in the figures.
